# An atypical NLR protein modulates the NRC immune receptor network

**DOI:** 10.1101/2021.11.15.468391

**Authors:** Hiroaki Adachi, Toshiyuki Sakai, Adeline Harant, Cian Duggan, Tolga O Bozkurt, Chih-hang Wu, Sophien Kamoun

**Affiliations:** The Sainsbury Laboratory, University of East Anglia, Norwich Research Park, NR4 7UH, Norwich, UK; Graduate School of Biological Sciences, Nara Institute of Science and Technology, Ikoma 630-0192, Japan; Laboratory of Crop Evolution, Graduate School of Agriculture, Kyoto University, Mozume, Muko, Kyoto 617-0001, Japan; Department of Life Sciences, Imperial College London, London, UK; Institute of Plant and Microbial Biology, Academia Sinica, Nankang, Taipei 11529, Taiwan

## Abstract

The NRC immune receptor network has evolved in asterid plants from a pair of linked genes into a genetically dispersed and phylogenetically structured network of sensor and helper NLR (nucleotide-binding domain and leucine-rich repeat-containing) proteins. In some species, such as the model plant *Nicotiana benthamiana* and other Solanaceae, the NRC network forms up to half of the NLRome, and NRCs are scattered throughout the genome in gene clusters of varying complexities. Here, we describe NRCX, an atypical, but essential member of the NRC family that lacks canonical features of these NLR helper proteins, such as a functional N-terminal MADA motif and the capacity to trigger autoimmunity. In contrast to other NRCs, systemic gene silencing of *NRCX* markedly impairs plant growth resulting in a dwarf phenotype. Remarkably, dwarfism of *NRCX* silenced plants is partially dependent on NRCX paralogs NRC2 and NRC3, but not NRC4. Despite its negative impact on plant growth when silenced systemically, transient RNA interference of *NRCX* in mature *N. benthamiana* leaves doesn’t result in visible cell death phenotypes. However, alteration of *NRCX* expression modulates the hypersensitive response mediated by NRC2 and NRC3 in a manner consistent with a negative role for NRCX in the NRC network. We conclude that NRCX is an atypical member of the NRC network that has evolved to contribute to the homeostasis of this genetically unlinked NLR network.

## INTRODUCTION

Plants are invaded by a multitude of pathogens and pests, some of which threaten food security in recurrent cycles of destructive epidemics. Yet, most plants are resistant to most parasites through their highly effective immune system. Plant defense consists of an expanded and diverse repertoire of immune receptors: cell-surface pattern recognition receptors (PRRs) and intracellular nucleotide-binding domain and leucine-rich repeat-containing (NLRs) proteins (Lu and Tsuda, 2021). Pathogen-associated molecular patterns (PAMPs) are recognized in the extracellular space by PRRs, resulting in pattern-triggered immunity (PTI) (Boutrot and Zipfel, 2017; DeFalco and Zipfel, 2021). NLRs perceive pathogen secreted proteins known as effectors and induce robust immune responses that generally include hypersensitive cell death (Jones et al., 2016; Kourelis and van der Hoorn, 2018; Saur et al., 2021). NLR-mediated immunity (also known as effector-triggered immunity) can be effective in restricting pathogen infection at invasion sites, and NLRs have also been recently shown to be involved in PRR-mediated signalling (Ngou et al., 2021a; Pruitt et al., 2021; Tian et al., 2021; Kourelis et al., 2021b). However, NLR-mediated immunity comes at a cost for plants. NLR mis-regulation and inappropriate activation can lead to deleterious physiological phenotypes, resulting in growth suppression and autoimmunity (Karasov et al., 2017), and the evolution of the plant immune system is constrained by fitness trade-offs between growth and immunity (Li and Weigel, 2021; Wan et al., 2021). However, our knowledge of the mechanisms by which diverse plant NLRs are regulated is still somewhat limited. Understanding how plants maintain NLR network homeostasis should help guide breeding disease resistant crops with limited fitness penalties.

NLRs occur across all kingdoms of life and generally function in innate immunity through “non-self” perception of invading pathogens (Jones et al., 2016; Uehling et al., 2017; Duxbury et al., 2021). Plant NLRs share a multidomain architecture typically consisting of a central NB-ARC (nucleotide-binding domain shared with APAF-1, various R-proteins and CED-4) and a C-terminal leucine-rich repeat (LRR) domain (Kourelis et al., 2021a). Plant NLRs can be sorted into sub-classes based on NB-ARC phylogenetic clustering and the type of N-terminal domain they carry (Shao et al., 2016; Kourelis et al., 2021a). The largest class of NLRs are the CC-NLRs with the Rx-type coiled-coil (CC) domain preceding the NB-ARC domain (Tamborski and Krasileva, 2020; Kourelis et al., 2021a; Lee et al. 2021). A prototypical ancient CC-NLR is the HOPZ-ACTIVATED RESISTANCE1 (ZAR1), which has remained relatively conserved throughout evolution over tens of millions of years (Adachi et al., 2021). However, the majority of CC-NLRs have massively expanded throughout their evolution, acquiring new activities through sequence diversification and integration of extraneous domains (Barragan and Weigel, 2021; Prigozhin and Krasileva, 2021), as well as losing particular molecular features following sub-functionalization (Adachi et al., 2019b).

Even though some plant NLRs function as singletons carrying both pathogen sensing and immune signalling activities, other NLRs form genetic and biochemical networks with varying degrees of complexity (Wu et al., 2018; Adachi et al., 2019a; Ngou et al., 2021b). NLRs can also cause deleterious genetic interactions known as hybrid incompatibility, presumably because mismatched NLRs inadvertently activate immunity (Li and Weigel, 2021; Wan et al., 2021). Hybrid autoimmunity is probably a trade-off of the rapidly evolving NLRome, which is expanding and diversifying in most angiosperm taxa, even at the intraspecific level (Van de Weyer et al., 2019; Lee and Chae, 2020; Prigozhin and Krasileva, 2021). NLRs can also cause a spontaneous autoimmune phenotype known as lesion mimicry (Bruggeman et al., 2015). Classic examples include mutants of the Toll/interleukin-1 receptor (TIR)-NLRs, *ssi4* (*suppressor of SA insensitivity 4*), *snc1* (*suppressor of npr1-1, constitutive 1*), *slh1* (*sensitive to low humidity 1*), *chs1* (*chilling-sensitive mutant 1*), *chs2* and *chs3*, which exhibit autoimmune phenotypes in Arabidopsis (Shirano et al., 2002; Zhang et al., 2003; Noutoshi et al., 2005; Huang et al., 2010; Yang et al., 2010; Bi et al., 2011; Wang et al., 2013). Other Arabidopsis NLRs, such as *laz5* (*lazarus 5*), *adr1* (*activated disease resistance 1*), *summ2* (*suppressor of mkk1 mkk2, 2*), *rps4* (*resistance to Pseudomonas syringae 4*), *csa1* (*constitutive shade-avoidance 1*), *soc3* (*suppressor of chs1-2, 3*), and *sikic* (*sidekick snc1*) are genetic suppressors of autoimmunity or cell death phenotypes (Palma et al., 2010; Bonardi et al., 2011; Zhang et al., 2012; Sohn et al., 2014; Xu et al., 2015; Zhang et al., 2017; Dong et al., 2018; Wu et al., 2020; Schulze et al., 2021). Nonetheless, to date, NLRs are not classed among so-called “lethal-phenotype genes” that are essential for plant viability and survival (Lloyd et al., 2015). This is despite the fact that ~500 *NLR* genes have been experimentally studied (Kourelis et al., 2021a).

Our understanding of molecular mechanisms underpinning plant NLR activation and the subsequent signalling events has significantly advanced with the elucidation of NLR protein structures. Activated CC-NLR ZAR1, TIR-NLRs RECOGNITION OF XOPQ 1 (ROQ1) and RESISTANCE TO PERONOSPORA PARASITICA 1 (RPP1) oligomerize into multimeric complexes known as resistosomes (Wang et al., 2019a; 2019b; Martin et al., 2020; Ma et al., 2020). In the case of ZAR1, activation by pathogens induces a switch from ADP to dATP/ATP binding and oligomerization into a pentameric resistosome (Wang et al., 2019b). This results in a conformational ‘death switch’, with the five N-terminal α1 helices forming a funnel-shaped structure that acts as a Ca^2+^ channel on the plasma membrane (Bi et al., 2021). The α1 helix of ZAR1 and about one-fifth of angiosperm CC-NLRs are defined by a molecular signature, the MADA motif (Adachi et al., 2019b). This α1/MADA helix is interchangeable between distantly related NLRs indicating that the ‘death switch’ mechanism applies to MADA-CC-NLRs from diverse plant taxa (Adachi et al., 2019b).

One class of MADA-CC-NLRs are the NRCs (NLR-REQUIRED FOR CELL DEATH), that are central nodes in a large NLR immune network of asterid plants (Wu et al., 2017). NRCs function as helper NLRs (NRC-H), required for a large number of sensor NLRs (NRC-S) to induce the hypersensitive response and immunity (Wu et al., 2017). These NRC-S are encoded by classical disease resistance genes that detect pathogens as diverse as viruses, bacteria, oomycetes, nematodes and insects. NRC-H and NRC-S form phylogenetic sister clades within a wider NRC superclade that makes up to half of the NLRome in the Solanaceae. This NRC superclade emerged early in asterid evolution about 100 million years ago from an ancestral pair of genetically linked NLRs (Wu et al., 2017). However, in sharp contrast to paired NLR pairs, functionally connected NRC-H and NRC-S genes are not always clustered and can be scattered throughout the plant genomes (Wu et al., 2017). Our current model is that the NRCs and their sensor evolved from bi-functional NLRs that have sub-functionalized and specialized throughout evolution (Wu et al., 2017; Adachi et al., 2019a; Adachi et al., 2019b). In support of this model, the MADA sequence has degenerated in NRC-S in contrast to the NRC-H, which carry functional N-terminal α1 helices (Adachi et al., 2019b).

Plant-pathogen coevolution has driven NLRs to form immune receptor networks (Wu et al. 2018; Ngou et al., 2021b). The emerging paradigm in the field of plant immunity is that helper NLRs, NRCs, as well as the CCR-NLRs ADR1 and N REQUIREMENT GENE 1 (NRG1), form receptor networks with multiple sensor NLRs (Adachi et al., 2019a; Feehan et al., 2020). These genetically dispersed NLR networks are likely to cause a heightened risk of autoimmunity during plant growth and development. Yet, the regulatory mechanisms that attenuate such deleterious effects of NLR networks are unknown. Here, we describe NRCX, an atypical NLR protein that belongs to the NRC-H phylogenetic clade. Gene silencing of *NRCX* markedly impairs plant growth, presumably because of mis-activation of its helper NLR paralogs NRC2 and NRC3. We propose that NRCX maintains NRC network homeostasis by modulating the activities of key helper NLR nodes during plant growth.

## RESULTS

### Systemic silencing of *NRCX* impairs *Nicotiana benthamiana* growth

Mis-regulated NLRs trigger physiological phenotypes in plants including a dwarfism associated with autoimmunity (Karasov et al., 2017; Li and Weigel, 2021; Wan et al., 2021). However, to date, NLRs have not been classed among the genes that are essential for plant viability (Lloyd et al., 2015). While performing virus-induced gene silencing (VIGS) of NRC family genes in the model plant *Nicotiana benthamiana* (*SI Appendix*, Fig. S1), we found that silencing of NLR NbS00030243g0001.1 (referred to from here on as NRCX) causes a severe dwarf phenotype (Fig. 1 A and B). In these VIGS experiments, we also independently targeted NRC helper (NRC-H) genes, *NRC2*, *NRC3* and *NRC4*, as well as the NRC sensor (NRC-S) gene *Prf* (NRC2/3 dependent) for silencing in *N. benthamiana* (Lu et al., 2003; Wu et al., 2017). Yet, none of the NRC-H and NRC-S silenced *N. benthamiana* plants showed quantitative growth defects (Fig. 1 A and B). These results suggest that *NRCX* is unique among NLRs described to date as a “lethal-phenotype” gene according to the definition of Lloyd et al. (2015). *NRCX* silencing could lead to NLR mis-regulation, thereby resulting in the dwarf phenotype in *N. benthamiana* plants.

**Fig. 1.**
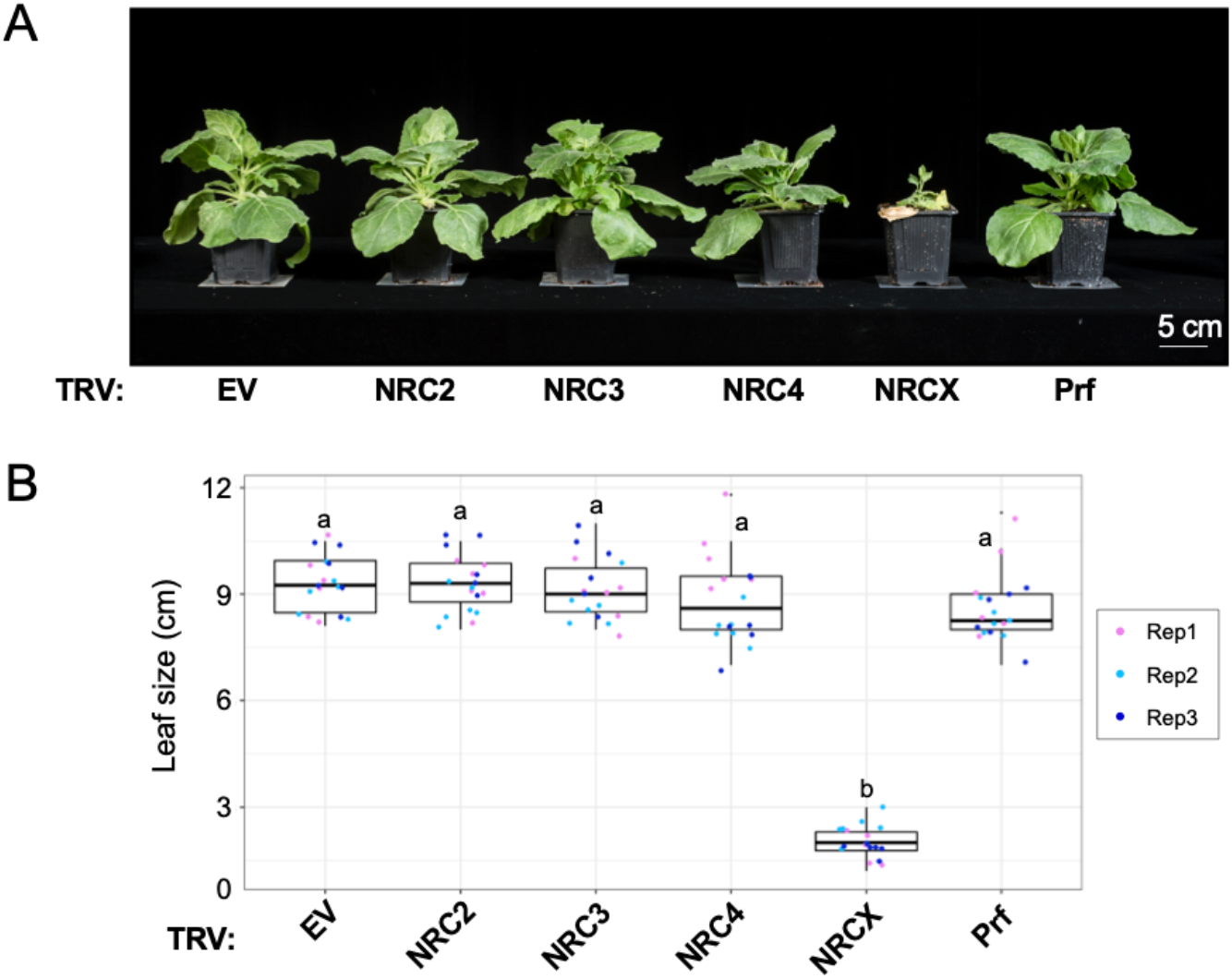
Virus-induced gene silencing of *NRCX* impairs *N. benthamiana* growth. **(A)** The morphology of 6-week-old *NRC2-*, *NRC3-*, *NRC4-*, *NRCX-* or *Prf*-silenced *N. benthamiana* plants. 2-week-old *N. benthamiana* plants were infiltrated with *Agrobacterium* strains carrying tobacco rattle virus (TRV) VIGS constructs, and photographs were taken 4 weeks after the agroinfiltration. TRV empty vector (TRV:EV) was used as a negative control. **(B)** Quantification of the leaf size of 6-week-old *NRC2-*, *NRC3-*, *NRC4-*, *NRCX-* or *Prf*-silenced *N. benthamiana* plants. One leaf per plant was harvested from the same position (the 5th leaf from cotyledons) and was used for measuring the leaf diameter. Statistical differences among the samples were analysed with Tukey’s HSD test (p<0.01).

### NRCX is a MADA-CC-NLR in the NRC helper phylogenetic clade

We investigated the precise phylogenetic position of NRCX in the NRC superclade. First, we built a phylogenetic tree with 431 CC-NLRs, including the CC-NLRome from four representative plant species (Arabidopsis, sugar beet, tomato and *N. benthamiana*) and 16 representative CC-NLRs (Fig. 2A). Next, we extracted the NRC-H subclade NLRs, which includes NRCX, for further phylogenetic analysis (Fig. 2 A and B). In this NRC-H subclade, NRCX forms a small clade together with a tomato NLR (Solyc03g005660.3.1; named as SlNRCX) (Fig. 2B, *SI Appendix*, Fig. S2). The NRCX clade is more closely related to NRC1/2/3 subclade than to the NRC4 subclade (Fig. 2B).

**Fig. 2.**
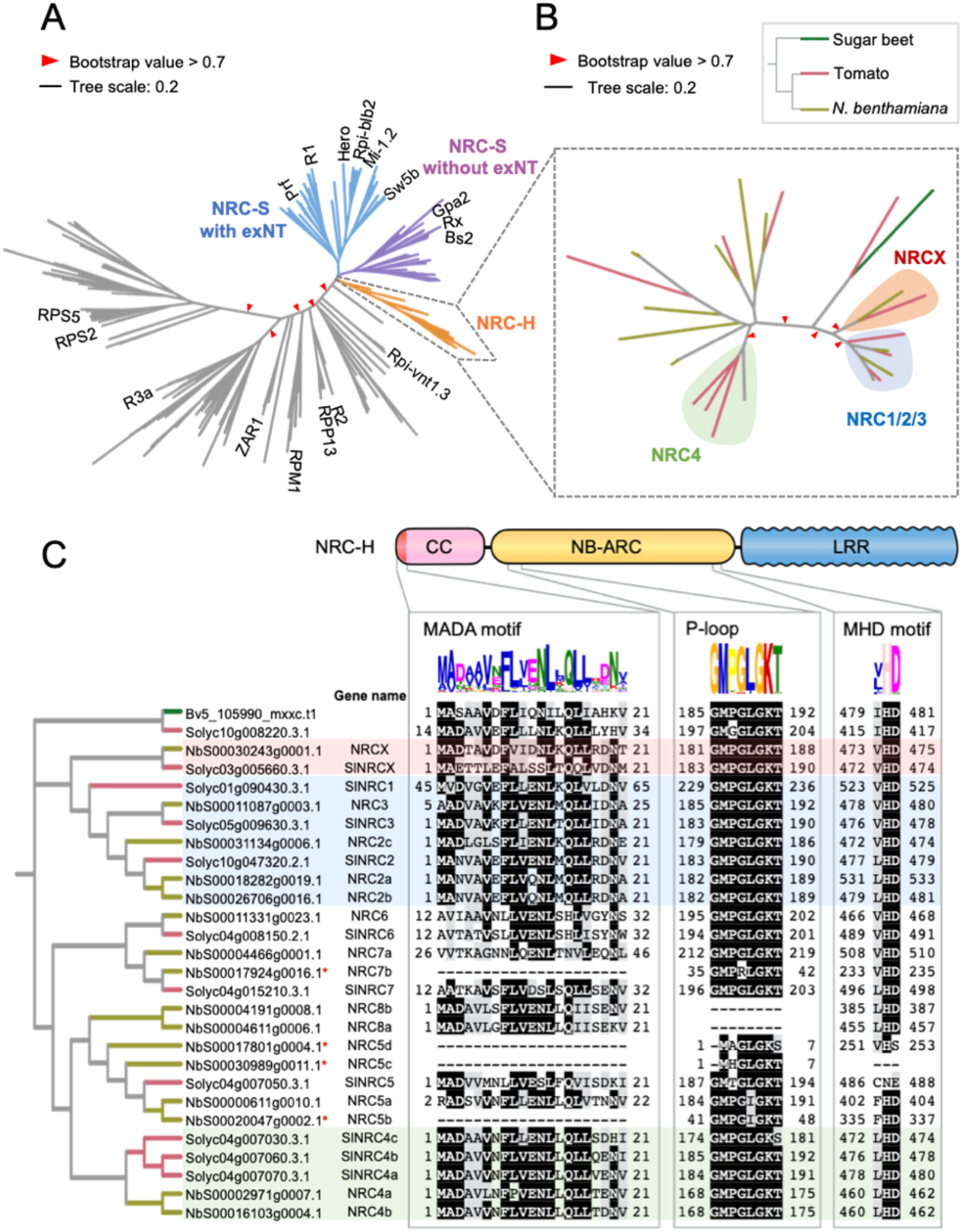
NRCX is a MADA-type CC-NLR in the NRC-helper clade. **(A, B)** NRCX is an NRC-helper (NRC-H) member phylogenetically closely related to the NRC1/2/3 clade. The maximum likelihood phylogenetic tree was generated in RAxML version 8.2.12 with JTT model using NB-ARC domain sequences of 431 NLRs identified from *N. benthamiana* (NbS-), tomato (Solyc-), sugar beet (Bv-), and Arabidopsis (AT-) (*SI Appendix*, File S3). The NRC superclade containing NRC-H and NRC-sensor (NRC-S) clades are described with different branch colours. The NRC-S clade is divided into NLRs that lack an extended N-terminal domain (exNT) prior to their CC domain and those that carry an exNT. The NRC-H clade phylogenetic tree is shown with different colours based on plant species (B). Red arrow heads indicate bootstrap support > 0.7 and is shown for the relevant nodes. The scale bars indicate the evolutionary distance in amino acid substitution per site. **(C)** Domain and motif architectures of NRC-H clade members. Amino acid sequences of MADA motif, P-loop and MHD motif are mapped onto the NRC-H phylogenetic tree. Each motif was identified in MEME using NRC-H sequences. NRCX, NRC1/2/3 and NRC4 clades are highlighted in red, blue and green, respectively.

We scanned NRCX and other NRC-H proteins for conserved sequence motifs (Fig. 2C). NRC-H members typically have the N-terminal MADA motif that is functionally conserved across many dicot and monocot CC-NLRs and is required for hypersensitive cell death and disease resistance (Adachi et al., 2019b). We ran the HMMER software (Eddy, 1998) to query NRCX with a previously reported MADA motif-Hidden Markov Model (HMM) (Adachi et al., 2019b). This HMMER search detected a MADA sequence at the N-terminus of NRCX (HMM score = 22.2) and in all other NRC-H except in four *N. benthamiana* NRC-H proteins that have N-terminal truncations (NbS00017924g0016.1, NbS00017801g0004.1, NbS00030989g0011.1 and NbS00020047g0002.1) (Fig. 2C). In addition, NRCX carries intact P-loop and MHD motif in its NB-ARC domain like the majority of CC-NLRs (Fig. 2C). We noted that the P-loop wasn’t predicted in NbS00004191g0008.1 and NbS00004611g0006.1, and the MHD motif is absent or divergent in NbS00030989g0011.1, NbS00017801g0004.1 and Solyc04g007050.3.1 (Fig. 2C). Taken together, we conclude that NRCX has the typical sequence motifs of MADA-CC-NLRs, similar to the great majority of proteins in the NRC-H subclade.

### The N-terminal MADA motif of NRCX is not functional and doesn’t mediate hypersensitive cell death

We investigated whether or not the MADA motif of NRCX is functional using the motif swap strategy we previously developed (Adachi et al., 2019b). To this end, we swapped the first 17 amino acids of an NRC4 autoactive MHD motif mutant (NRC4^DV^) with the equivalent region of NRCX, resulting in a MADA^NRCX^-NRC4^DV^ chimeric protein (Fig. 3A-B). The N-terminal sequence of NRC4 can be functionally replaced with matching sequences of other MADA-CC-NLRs, including NRC2 from *N. benthamiana*, ZAR1 from Arabidopsis and even the monocot MADA-CC-NLRs MLA10 and Pik-2 (Adachi et al., 2019b). Intriguingly, MADA^NRCX^-NRC4^DV^ did not induce any visible cell death when expressed by agroinfiltration in *N. benthamiana* leaves unlike the positive controls MADA^NRC2^-NRC4^DV^ and MADA^ZAR1^-NRC4^DV^ (Fig. 3A-C). MADA^NRCX^-NRC4^DV^ chimeric protein accumulated to similar levels as MADA^ZAR1^-NRC4^DV^ in *N. benthamiana* leaves, indicating that the lack of activity was probably not due to protein destabilization (Fig. 3E). These results indicate that the MADA region of NRCX does not have the capacity to trigger hypersensitive cell death in the NRC4 protein background, and therefore despite its sequence conservation probably forms a non-functional N-terminal α1 helix.

**Fig. 3.**
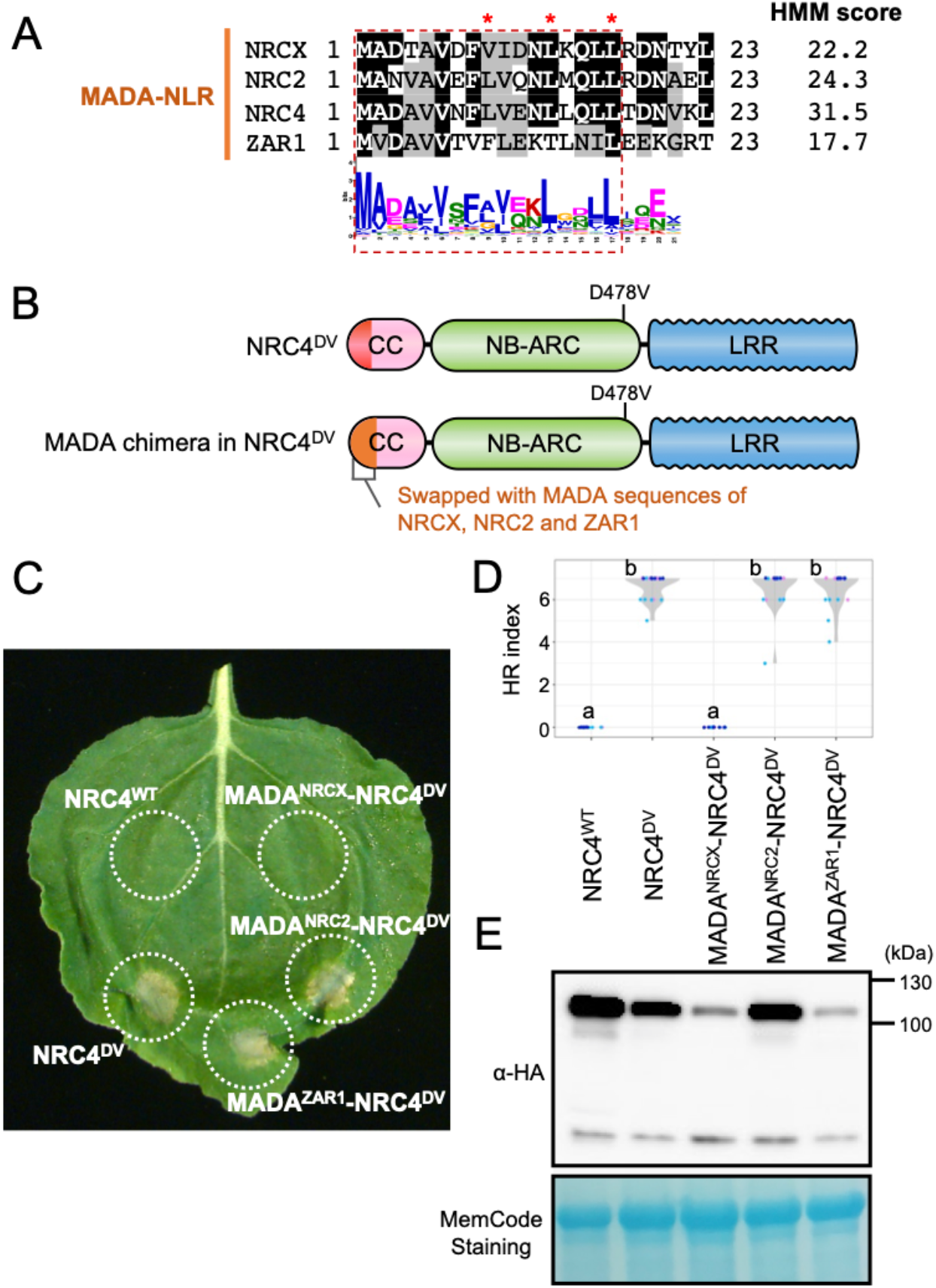
Unlike other MADA-CC-NLRs, the N-terminal 17 amino acids of NRCX fails to confer cell death activity to an NRC4 autoactive mutant. **(A)** Alignment of the N-terminal region of the MADA-CC-NLRs, NRCX, NRC2, NRC4 and ZAR1. Key residues for cell death activity (Adachi et al. 2019b) are marked with red asterisks in the sequence alignment. Each HMM score is indicated. **(B)** Schematic representation of NRC4 MADA motif chimeras. The first 17 amino acid region of NRCX, NRC2 and ZAR1 was swapped into the autoactive NRC4 mutant (NRC4^DV^), resulting in the NRC4 chimeras with MADA sequences originated from other MADA-CC-NLRs. **(C)** Cell death phenotypes induced by the NRC4 chimeras. NRC4^WT^-6xHA, NRC4^DV^-6xHA and the chimeras were expressed in *N. benthamiana* leaves by agroinfiltration. Photographs were taken at 5 days after the agroinfiltration. **(D)** Violin plots showing cell death intensity scored as an HR index based on three independent experiments. Statistical differences among the samples were analyzed with Tukey’s honest significance difference (HSD) test (p<0.01). **(E)** *In planta* accumulation of the NRC4 variants. For anti-HA immunoblots of NRC4 and the mutant proteins, total proteins were prepared from *N. benthamiana* leaves at 1 day after the agroinfiltration. Equal loading was checked with Reversible Protein Stain Kit (Thermo Fisher).

### Unlike other NRC helpers, mutations in the MHD motif fail to confer autoactivity to NRCX

The absence of a functional MADA/α1 helix in NRCX hints at the possibility that this NLR lacks a cell death-inducing activity. To challenge this hypothesis, we performed site-directed mutagenesis of the MHD motif predicted in the NB-ARC domain of NRCX. Previously, we showed that histidine (H) to arginine (R) or aspartic acid (D) to valine (V) substitutions within the MHD motif confer autoactivity to the NRC helpers NRC2, NRC3 and NRC4 (Derevnina et al., 2021). Therefore, we first introduced the HR and DV mutations in NRCX (Fig. 4A). Both NRCX^HR^ and NRCX^DV^ did not induce macroscopic cell death when expressed in *N. benthamiana* leaves using agroinfiltration, in contrast to the autoactive NRC4^DV^ (Fig. 4B). To further challenge this finding, we randomly mutated NRCX H474 and D475 residues in the MHD motif and used agroinfiltration to express them in *N. benthamiana* leaves (*SI Appendix*, Fig. S3). None of the 55 independent NRCX MHD mutants we tested induced visible macroscopic cell death (*SI Appendix*, Fig. S3). These results indicate that NRCX does not have the capacity to induce cell death, unlike the other NRC-H it is related to.

**Fig. 4.**
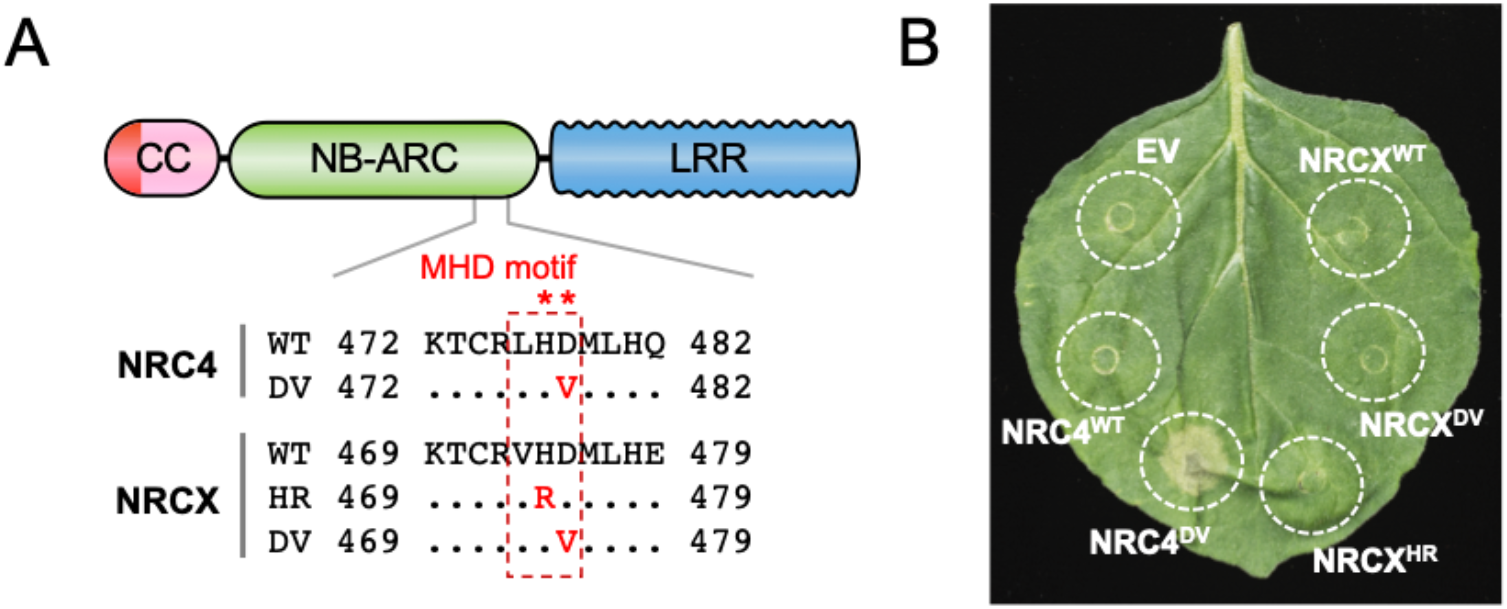
Mutations in the NRCX MHD motif do not result in autoactive cell death in *N. benthamiana*. **(A)** Schematic representation of the mutated sites in the NRC4 and NRCX MHD motifs. Substituted residues are shown in red in the multiple sequence alignment. **(B)** NRC4^WT^, NRCX^WT^ and the MHD mutants were expressed in *N. benthamiana* leaves by agroinfiltration. Cell death phenotype induced by the MHD mutant was recorded 5 days after the agroinfiltration. Quantification of the cell death intensity is shown in *SI Appendix*, Fig. S3.

### The dwarf phenotype of *NRCX*-silenced *Nicotiana benthamiana* plants is partially dependent on NRC2 and NRC3 but not NRC4

We hypothesized that NRCX has a regulatory role in the NRC network, given that it belongs to the NRC helper clade but lacks the capacity to cause hypersensitive cell death. To test this hypothesis, we silenced *NRCX* in three different *N. benthamiana* lines, *nrc2/3*, *nrc4* and *nrc2/3/4*, that we previously described as carrying loss-of-function mutations in NRC2, NRC3 and/or NRC4 (Adachi et al., 2019b; Wu et al., 2020; Witek et al., 2021) (Fig. 5). Four weeks after inoculation with TRV:*NRCX*, we observed partial suppression of the TRV:*NRCX*-mediated growth defects in *nrc2/3* and *nrc2/3/4* plants, but not in *nrc4* plants (Fig. 5 A-C). We confirmed that the quantitative differences were reproducible by using at least two independent lines of each of the three mutants (Fig. 5 A-C). In these experiments, we did not observe quantitative growth differences between *nrc2/3* and *nrc2/3/4* plants (Fig. 5 B and C).

**Fig. 5.**
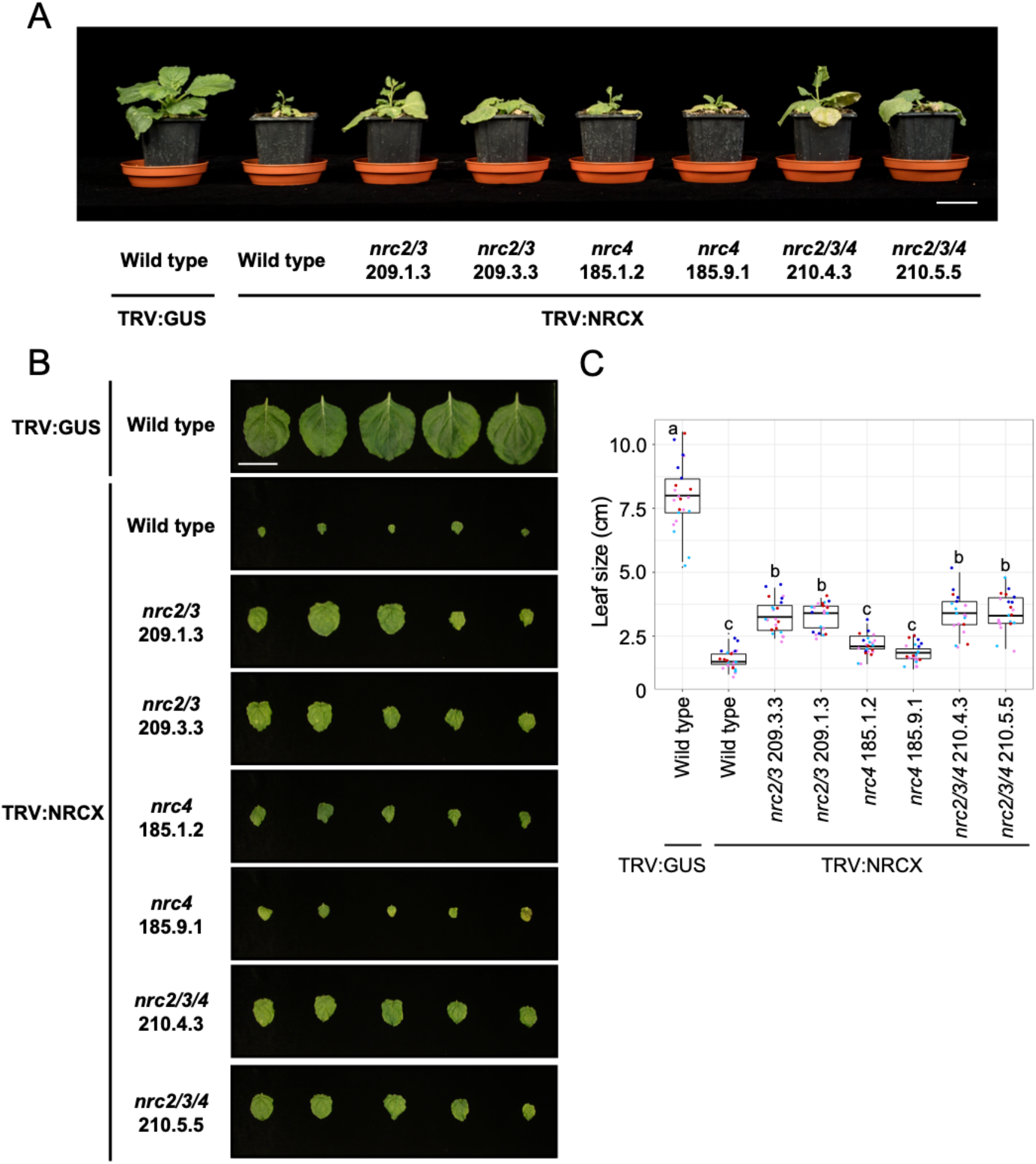
The dwarf phenotype by *NRCX*-silenced *N.* benthamiana plants is partially dependent on NRC2 and NRC3 but not NRC4. **(A)** The morphology of 6-week-old wild-type *N. benthamiana*, *nrc2/3*, *nrc4* and *nrc2/3/4* CRISPR-knockout lines expressing TRV:NRCX. 2-week-old wild-type and the knockout plants were infiltrated with *Agrobacterium* strains carrying TRV:GUS or TRV:NRCX, and photographs were taken 4 weeks after the agroinfiltration. **(B, C)** Quantification of the leaf size. One leaf per each plant was harvested from the same position (the 5th leaf from cotyledons) and was used for measuring the leaf diameter. Statistical differences among the samples were analyzed with Tukey’s HSD test (p<0.01). Scale bars = 5 cm.

To independently challenge these results, we performed co-silencing experiments where *NRCX* was knocked-down together with *NRC2/3*, *NRC4* or *NRC2/3/4*. To silence multiple genes by VIGS in *N. benthamiana* plants, we tandemly cloned gene fragments of *NRCX* and other NRCs in the same TRV expression vector. Compared to TRV:*NRCX* plants, the dwarf phenotype was partially suppressed in TRV:*NRC2/3/X* and TRV:*NRC2/3/4/X*, but not in TRV:*NRC4/X* plants (*SI Appendix*, Fig. S4 A-C). Each silencing construct specifically reduced mRNA levels of the target *NRC* gene, indicating that phenotypic differences are unlikely to be due to off-target gene silencing effects (*SI Appendix*, Fig. S4D). Altogether, we conclude that the TRV:*NRCX*-mediated dwarf phenotype is partially dependent on NRC2 and NRC3, but not NRC4.

### RNA interference of *NRCX* in *Nicotiana benthamiana* leaves doesn’t result in visible cell death phenotypes

To gain further insights into NRCX activities, we generated a hairpin-silencing construct (RNAi:*NRCX*) for *NRCX* silencing after transient expression in mature *N. benthamiana* leaves. First, we investigated the degree to which the RNAi:*NRCX* silencing construct causes cell death in *N. benthamiana* leaves, given that NLR-mediated dwarfism in plants is often linked to cell death (Zhang et al., 2012). Five days after the agroinfiltration, RNAi:*NRCX* did not result in macroscopic cell death in *N. benthamiana* leaves whereas co-expression of Pto/AvrPto resulted in a visible cell death response (*SI Appendix*, Fig. S5A). To monitor cell death at a microscopic level, we performed trypan blue staining, which generally visualizes dead cells. Trypan blue only stained trichomes in the RNAi:*NRCX* and RNAi:*GUS* (negative control) treated leaf panels, whereas it clearly revealed cell death in epidermal and mesophyll cells in the Pto/AvrPto treatment (positive control) (*SI Appendix*, Fig. S5 B and C). These observations indicate that RNA interference of *NRCX* does not cause visible cell death in *N. benthamiana* leaves. This RNAi strategy also enabled us to bypass the severe dwarf phenotype observed in whole-plant VIGS experiments to perform functional analyses of NRCX.

### RNA interference of *NRCX* enhances NRC2- and NRC3-dependent cell death

Our finding that NRC2 and NRC3 are genetic suppressors of NRCX raises the possibility that NRCX negatively modulates the activity of these helper NLRs and prompted us to investigate this hypothesis. To this end, we co-expressed by agroinfiltration in *N. benthamiana* leaves RNAi:*NRCX* with NRC-dependent sensor NLRs (Pto/SlPrf, Gpa2, Rpi-blb2, R1, Sw-5b and Rx) or NRC-independent NLRs (R2 and R3a) and their matching pathogen effectors (Wu et al., 2017; Derevnina et al., 2021). *NRCX* silencing by the hairpin construct enhanced the hypersensitive cell death triggered by the sensor NLRs SlPrf, Gpa2 (NRC2 and NRC3 dependent) and Sw-5b (NRC2, NRC3 and NRC4 dependent) relative to the RNAi:*GUS* control, but did not affect the other sensor NLRs, including Rpi-blb2, R1, Rx, R2 and R3a (Fig. 6 A and 6B and *SI Appendix*, Fig. S6). It should be pointed out that although NRC2, NRC3 and NRC4 redundantly contribute to effector-activated Sw-5b hypersensitive cell death (Wu et al., 2017), we recently showed that an autoactive mutant of Sw-5b (Sw-5b^D857V^) signals only through NRC2 and NRC3 (Derevnina et al., 2021). Therefore, we conclude that knocking-down of *NRCX* enhances the activities of NRC2- and NRC3-dependent NRC-S, but doesn’t affect the activity of NRC-S that are only dependent on NRC4, as well as other NLR outside the NRC network.

**Fig. 6.**
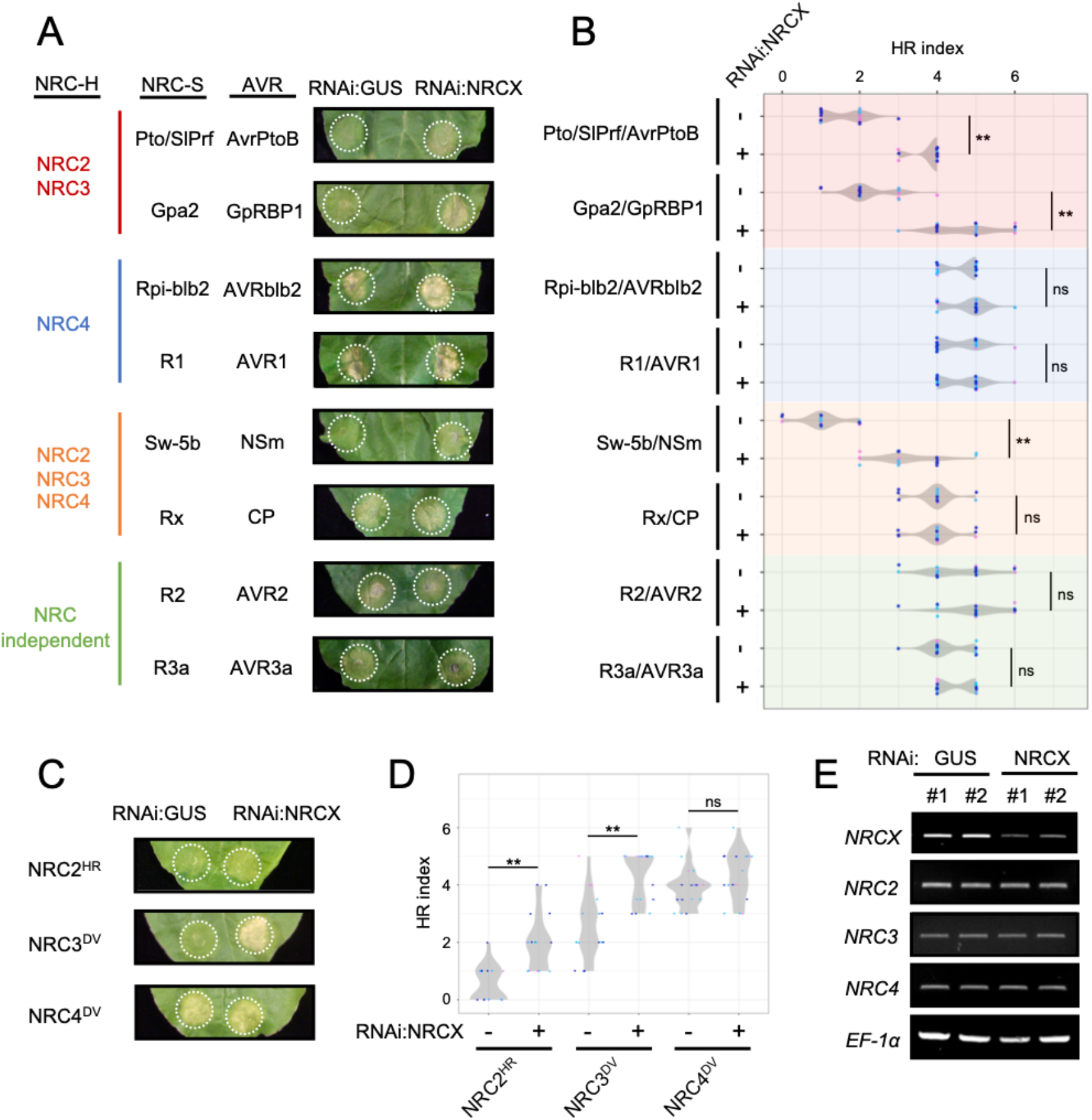
RNA interference of *NRCX* enhances NRC2- and NRC3-dependent hypersensitive cell death. **(A)** Hypersensitive cell death phenotypes after co-expressing different NRC-S and AVR combinations with RNAi:GUS (control) or RNAi:NRCX by agroinfiltration. Cell death intensity was scored at 2-5 days after the agroinfiltration, and photographs were taken at 5 days after the agroinfiltration. **(B)** Violin plots showing cell death intensity scored as an HR index at 5 days after the agroinfiltration. Time-lapse HR index is shown in *SI Appendix*, Fig. S6. The HR index plots are based on three independent experiments. Asterisks indicate statistically significant differences with *t* test (**p<0.01). **(C)** Autoactive cell death phenotypes induced by MHD mutants of NRC2, NRC3 and NRC4 with RNAi:GUS or RNAi:NRCX. Photos indicate cell death response at 4 days after the agroinfiltration. **(D)** Violin plots showing cell death intensity scored at 4 days after the agroinfiltration. Time-lapse HR index is shown in *SI Appendix*, Fig. S7. **(E)** *NRCX* silencing in *N. benthamiana*. Leaf samples were collected 2 days after agroinfiltration expressing RNAi:GUS and RNAi:NRCX. Total RNA was extracted from two independent plant samples (#1, #2). The expression of NRCX and other NRC-H genes were analysed in semi-quantitative RT-PCR using specific primer sets. Elongation factor 1α (EF-1α) was used as an internal control.

Next, we investigated the extent to which NRCX affects the activities of autoactive mutants of the NRC-H, NRC2^H480R^ (NRC2^HR^), NRC3^D480V^ (NRC3^DV^) and NRC4^D478V^ (NRC4^DV^), which cause cell death even in the absence of pathogen effectors (Derevnina et al., 2021). We co-expressed NRC2^HR^, NRC3^DV^ and NRC4^DV^ with RNAi:*NRCX* or RNAi:*GUS* by agroinfiltration in *N. benthamiana* leaves. *NRCX* silencing enhanced the cell death responses caused by NRC2^HR^ and NRC3^DV^, but not NRC4^DV^, compared to the RNAi:*GUS* silencing control (Fig. 6 C and D and *SI Appendix*, Fig. S7). RNAi:*NRCX* expression reduced mRNA levels of endogenous *NRCX* gene, but did not alter the expression of other NRCs, indicating that the enhanced cell death phenotype was probably not due to off-target silencing (Fig. 6E). Altogether, these two sets of RNAi experiments indicate that NRCX modulates the helper NLRs NRC2 and NRC3 nodes in the NRC network.

### NRCX overexpression compromises NRC2 and NRC3, but not NRC4, autoimmune cell death

We further challenged our model by testing the extent to which NRCX overexpression can suppress the cell death caused by autoactive NRC2 and NRC3. We generated an overexpression construct of wild-type NRCX and co-expressed it by agroinfiltration with autoactive NRC2^HR^, NRC3^DV^ and NRC4^DV^ mutants in *N. benthamiana* leaves. Five days after agroinfiltration, we observed that wild-type NRCX expression compromised autoimmune cell death triggered by NRC2^HR^ and NRC3^DV^, but not NRC4^DV^, relative to the empty vector control (*SI Appendix*, Fig. S8). Taken together, manipulation of *NRCX* expression levels in *N. benthamiana* leaves suggests a negative role of NRCX in NRC2 and NRC3, but not NRC4, mediated immunity.

### *NRCX* is differentially expressed relative to *NRC2a/b* following activation of pattern-triggered immunity

Our finding that NRCX modulates NRC2 and NRC3 nodes prompted us to investigate the transcriptome dynamics of *NRCX* compared to these *NRC-H* in different plant tissues. To obtain transcriptome data of NRC-H clade members, we extracted total RNA from leaf, root and flower/bud of six-week-old wild-type *N. benthamiana*. Three replicate each from independent plants were subjected to RNA-seq and resulted in 40 million 150-bp paired-end reads per sample. Then, we calculated Transcripts Per Million (TPM) values for *N. benthamiana NLR* genes (*SI Appendix*, Table S1). In Fig. 7a, we focused on transcriptome profiles of NRC-H with a TPM > 1.0 and found that *NRCX*, *NRC2a/b/c*, *NRC3* and *NRC4a/b* genes were expressed in all three tissues, leaf, root and flower/bud at relatively similar ratios. Notably, *NRCX* was about 3 to 5-fold more highly expressed in roots (TPM = 25.5) compared to leaves (TPM = 7.5) and flowers/buds (TPM = 5.2) (Fig. 7a). In addition to the *NRC2*, *NRC3*, *NRC4* and *NRCX*, we also noted that *NRC5a* and *NRC8a* were expressed in the three tissues, whereas *NRC7a* and *NRC8b* were mainly expressed in *N. benthamiana* roots (Fig. 7a). These transcriptome profiles suggest that NRCX maintains NRC2/3 subnetwork homeostasis through plant growth and development in leaf, root and flower/bud.

**Fig. 7.**
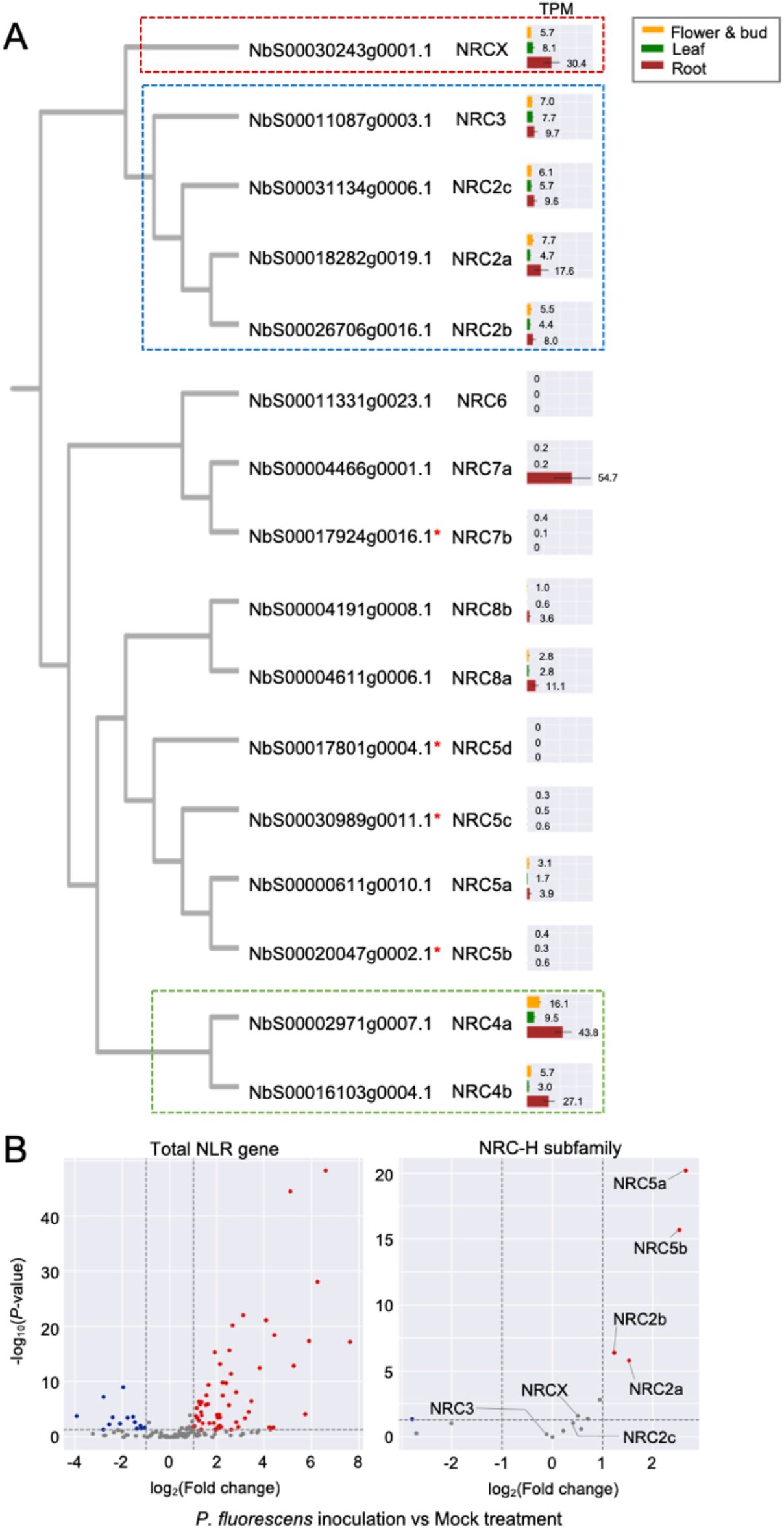
*NRCX* is differentially expressed relative to *NRC*2*a/b* genes in *N. benth* amiana leaves after *Pseudomonas fluorescens* inoculation. **(A, B)** TPM values were calculated using RNA-seq data of three different tissues (leaf, root, flower and bud) in 6-weeks old *N. benthamiana* plants and published transcriptome data of *N. benthamiana* leaves with mock treatment or *P. fluorescens* inoculation (Pombo et al., 2019). **(A)** The TPM values analysed from the three different tissue samples are mapped onto phylogeny extracted from the phylogenetic tree in Fig 2C. Red asterisks indicate truncated genes. **(B)** Volcano plots show up-regulated genes (red dots: log_2_(*P. fluorescens*/mock) ≧ 1 and *P*-value ≦ 0.05) and down-regulated genes (blue dots: log_2_(*P. fluorescens*/mock) ≦ −1 and *P*-value ≦ 0.05) in response to *P. fluorescens* inoculation compared to mock treatment.

Next, we analysed transcriptome data of six-week-old wild-type *N. benthamiana* leaves upon inoculation with avirulent bacteria *Pseudomonas fluorescens* (Pombo et al., 2019). *P. fluorescens* inoculation is considered to trigger PTI in *N. benthamiana*. We, therefore, explored the degree to which *NRC-H* genes are up- or down-regulated upon PTI activation. In total, 61 *NLR* genes were up-regulated, while 14 *NLR* genes were down-regulated in bacterial vs. mock treatment (Fig. 7b). Interestingly, the expression of *NRC2a/b* genes was up-regulated in response to *P. fluorescens* inoculation whereas there were no particular differential expression changes with *NRCX* (Fig. 7b; *SI Appendix*, Table S2). Other *NRC-H* genes, such as *NRC2c*, *NRC3* and *NRC4a*/*b*, were also unchanged whereas the expression level of *NRC5a/b* increased following bacterial treatment (Fig. 7b). We conclude that following activation of immunity in response to bacterial inoculation, *NRC2a/b* become more highly expressed than their paralog and modulator *NRCX* thereby altering the homeostasis of the NRC network.

## DISCUSSION

Co-operating plant NLRs are currently categorized into “sensor NLR” for effector recognition or “helper NLR” for immune signalling (Adachi et al., 2019a; Feehan et al., 2020). These functionally specialized NLR sensors and helpers function in pairs or networks across many species of flowering plants. In this study, we found that NRCX, a recently diverged paralog of the NRC class of helper NLRs, is essential to sustain proper plant growth and is a modulator of the genetically dispersed NRC2/3 subnetwork. We propose that NRCX has evolved to maintain the homeostasis of at least a section of the NRC network (Fig. 8). NRCX is also atypical as far as NLR helpers and NRC proteins go, lacking a functional N-terminal MADA motif and the capacity to trigger hypersensitive cell death. At the moment, we cannot categorize NRCX as either a sensor or helper NLR, and it is best described as an NLR modulator.

**Fig. 8.**
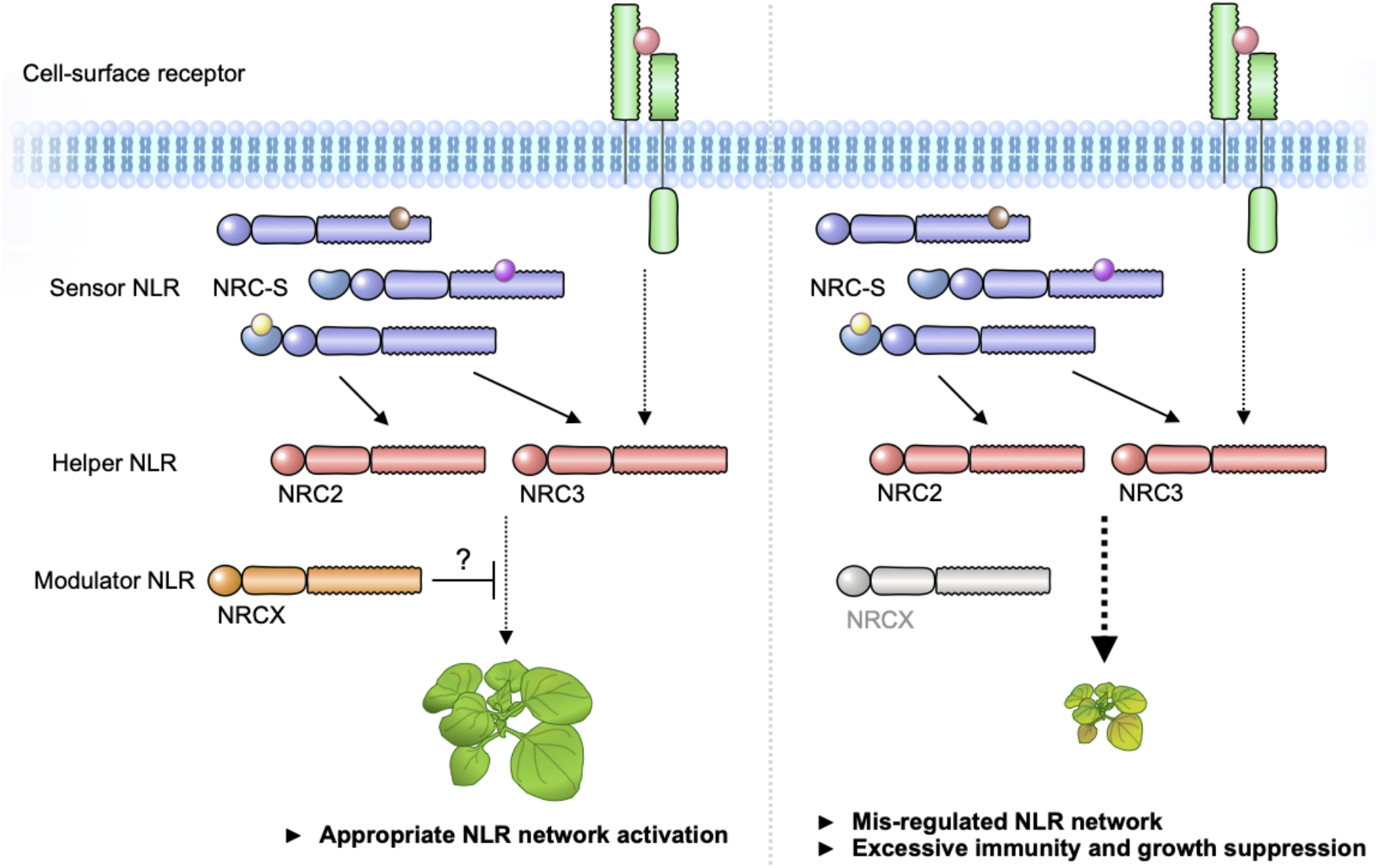
Modulator NLR has evolved to maintain NLR network homeostasis. We propose that “Modulator NLR” contributes to NLR immune receptor network homeostasis during plant growth. A modulator NRCX has a similar sequence signature with helper MADA-CC-NLRs, but unlike helpers, NRCX lacks the functional MADA motif to execute cell response. NRCX modulates the NRC2/NRC3 subnetwork composed of multiple sensor NLRs and cell-surface receptor (left). Loss of function of NRCX leads to the enhanced hypersensitive response and dwarfism in *N. benthamiana* plants (right).

NLRs are often implicated in spontaneous autoimmune phenotypes in plants and humans (Karasov et al., 2017; Feerick and McKernan, 2017). In plants, autoimmunity is often observed at the F1 and later generations when genetically distinct plant accessions are crossed (Karasov et al., 2017; Li and Weigel, 2021; Wan et al., 2021). One well-studied case of hybrid autoimmunity is induced by a hetero-complex of NLRs from genetically unlinked *NLR* gene loci between DANGEROUS MIX 1 (DM1) from Arabidopsis accession Uk-3 and DM2 from Uk-1 (Bomblies et al., 2007; Chae et al., 2014; Tran et al., 2017). Therefore, hybrid incompatibility can be due to inappropriate activation of mismatched NLRs. Amino acid insertions or substitutions in *NLR* genes can also exhibit autoimmune phenotypes in Arabidopsis, such as in the NLR alleles *ssi4* (G422R), *snc1* (E552K), *slh1* (single leucine insertion to RRS1 WRKY domain), *chs1* (A10T), *chs2* (S389F) and *chs3-2D* (C1340Y) (Shirano et al., 2002; Zhang et al., 2003; Noutoshi et al., 2005; Huang et al., 2010; Bi et al., 2011; Wang et al., 2013). In *chs3-1*, a truncation of the C-terminal LIM-containing domain causes autoimmunity (Yang et al., 2010). However, to date, T-DNA insertion or deletion mutants including *chs3-2*, where expression of full-length NLR modules is suppressed, do not show autoimmune phenotypes (Zhang et al., 2003; Yang et al., 2010; Narusaka et al., 2016; Zhang et al., 2017). Therefore, autoimmune *NLR* mutants are typically considered gain-of-function mutants. A striking finding in this study is that *NRCX* silencing resulted in severe dwarfism (Fig. 1). To our knowledge, this finding is a unique example where loss-of-function of an *NLR* gene causes a “lethal” plant phenotype, following the definition by Lloyd et al. (2015). Therefore, *NRCX* can be classified as an essential plant gene, which is exceptional given that *NLR* genes tend to act just the opposite way, by causing fitness penalties.

NRC2 and NRC3 partially suppress the dwarf phenotype of *NRCX*-silenced plants and can be viewed as genetic suppressors of NRCX (Fig. 1 and Fig. 5). These findings inspired us to draw a model which expands our understanding of the NRC network to include NRCX as a modulator of the NRC2 and NRC3 nodes (Fig. 8). This network even connects NLRs to cell-surface immune receptors, given that Kourelis et al. (2021b) recently showed that NRC3 is required for the hypersensitive cell death triggered by the receptor protein Cf-4. We propose that NRCX contributes to the homeostasis of the NRC2/3 subnetwork, which are hubs in an immune network composed of both intracellular NRC-S and the cell-surface receptors. When the expression level of *NRCX* is reduced, NRC2 and NRC3 cause autoimmunity, possibly through inadvertent activation (Fig. 8). However, transient RNA interference of *NRCX* in *N. benthamiana* leaves did not cause autoimmune cell death (*SI Appendix*, Fig. S5), although it enhanced the hypersensitive cell death elicited by effector recognition (Fig. 6). This indicates that NRCX-mediated dwarfism is trigger-dependent and may occur in particular tissues and/or during certain developmental stages presumably in a manner similar to pathogen recognition. Considering that *nrc2/nrc3* knockout plants do not fully recover from dwarfism (Fig. 5), it is possible that *NRCX* silencing leads to a permanent “trigger-happy” status of NRC2/3 and other NLR(s) that ultimately perturbs plant growth.

NRCX clusters within the NRC-H subclade, which is populated with proteins with a MADA-CC-NLR architecture (Fig. 2). However, the N-terminal MADA motif of NRCX was not functional when swapped into NRC4 (Fig. 3), compared to other similar swaps we previously tested (Adachi et al., 2019b; Duggan et al., 2021a; Kourelis et al. 2021b). We conclude that the MADA motif of NRCX has degenerated from the canonical sequence present in the typical NRC helper NLRs and lost the capacity to execute the hypersensitive cell death response. In fact, a polymorphism in the NRCX MADA motif region, Thr-4, is unique to this protein among NRC-H (Fig. 2). Therefore, loss of cell death executor activity in NRCX might represent another path in the evolution of networked NLRs besides sensor and helper NLRs, as in the NLR functional specialization “use-it-or-lose-it” model described Adachi et al. (2019b) and Kamoun (2021). We propose that NRCX has also functionally specialized into an atypical NLR protein that operates as an NLR modulator.

NRCX overexpression compromised but did not fully suppress NRC2 and NRC3 autoactive cell death (*SI Appendix*, Fig. S8). Its suppressor activity contrasts with that of the SPRYSEC15 effector from the cyst nematode *Globodera rostochiensis* and AVRcap1b from the oomycete *Phytophthora infestans*, which strongly suppress the cell death inducing activity of autoimmune NRC2 and NRC3 (Derevnina et al., 2021). In the future, it will be interesting to determine the mechanism by which NRCX modulates the activities of NRC-H proteins, and how that process compares with pathogen suppression of the NRC network.

Our work on NRCX adds to a growing body of knowledge about NLRs that modulate the activities of other NLRs. Classic examples include genetically linked NLR pairs, such as the Pia RGA5/RGA4 pair, in which the sensor RGA5 suppresses the RGA4 autoimmune cell death observed in *N. benthamiana* (Césari et al., 2014). In this and other one-to-one paired NLRs, the sensor NLR carries a regulatory role of its genetically linked helper mate that is released by the matching pathogen effector. In contrast, NRCX modulates a genetically dispersed NLR network composed of a large number of sensor NLRs and two helper NLRs, NRC2 and NRC3. Recently, Wu et al. (2021) showed that overexpression of NRG1C, antagonizes autoimmunity by its paralog NRG1A, *chs3-2D* and *snc1*, without affecting *chs1*, *chs2* autoimmunity and RPS2- and RPS4-mediated immunity. NRG1 is a member of CCR-NLR helper subfamily that triggers cell death via its N-terminal CCR domain (Jacob et al., 2021). Intriguingly, unlike NRCX, NRG1C has a truncated NLR architecture, lacking the whole N-terminal CCR domain and is therefore unlikely to execute the hypersensitive cell death (Wu et al., 2021). The emerging picture is that NRCX and NRG1C are an atypical subclass of helper NLRs, we term modulator NLRs, that lost their cell death executor activity through regressive evolution and evolved to modulate the activities of multiple NLR helper proteins. These examples form another case of functional specialization during the transitions associated with NLR evolution (Kamoun, 2021). We need a better appreciation of the diversity of structures and functions that come with the protein we classify as NLR immune receptors and integrate the different ways NLRs contribute to immunity (Adachi et al., 2019a; Feehan et al., 2020; Kourelis et al. 2021a; Kamoun, 2021).

Plant NLRs are tightly regulated at the transcript level because increased expression of NLRs can result in autoimmune phenotypes and fitness costs (Li et al., 2007; Gloggnitzer et al., 2014; Tsuchiya and Eulgem, 2013; Lai et al., 2020). However, activation of plant defense is associated with massive up-regulation of *NLR* genes. For instance, in Arabidopsis, dozens of *NLR* genes are up-regulated in response to pathogen-related treatments, such as the bacterial PAMP flg22 (Mohr et al., 2010; Yu et al., 2013). The current model is that NLR expression level is maintained at a low basal level but is amplified in the cells upon activation of pathogen-induced immunity. In our analyses, we found that 61 *NLR* genes, including *NRC2a/b*, are up-regulated in *N. benthamiana* leaves following *P. fluorescens* inoculation, while *NRCX* expression levels remain unchanged (Fig. 7b). This marked shift in the balance between *NRC2a/b* to *NRCX* expression levels may potentiate and amplify the activity of the NRC network resulting in more robust immune responses (Fig. 8).

The NRC superclade forms a large and complex NLR immune receptor network in asterid plants that connects to cell surface receptors (Wu et al., 2017; Kourelis et al. 2021b). Here we show that NRCX modulates the hub NLR proteins NRC2 and NRC3, but doesn’t affect their paralog NRC4. Further studies are required to determine the molecular mechanisms underpinning NRCX antagonism of NRC2- and NRC3-mediated immune response and what other mechanisms modulate other sections of the network, such as the NRC4 subnetwork. Our findings also highlight the potential fitness costs associated with expanded NLR networks as the risk of inadvertent activation increases with the network complexity. Indeed, we found that NRCX regulation of NRC2 and NRC3 is essential for normal plant growth and development. Further understanding of how NLR network homeostasis is maintained will provide insights for future breeding of vigorous and disease resistant crops.

## MATERIALS AND METHODS

### Plant growth conditions

Wild-type and mutant *Nicotiana benthamiana* were propagated in a glasshouse and, for most experiments, were grown in a controlled growth chamber with temperature 22–25 °C, humidity 45–65% and 16/8 hr light/dark cycle. The NRC knockout lines used have been previously described: *nrc2/3*-209.1.3 and *nrc2/3*-209.3.3 (Witek et al., 2021), *nrc4*-185.1.2 and *nrc4-*185.9.1 (Adachi, et al., 2019b), and *nrc2/3/4*-210.4.3 and *nrc2/3/4*-210.5.5 (Wu et al., 2020).

### Plasmid constructions

The cDNA of *NRCX* was amplified by PCR from *N. benthamiana* cDNA using Phusion High-Fidelity DNA Polymerase (Thermo Fisher) and primers listed in *SI Appendix*, Table S3. The PCR product was cloned into pICSL01005 (Level 0 acceptor for CDS no stop modules, Addgene no.47996) as a level 0 module for Golden Gate assembly. To generate NRCX-3xFLAG overexpression construct, the pICSL01005::NRCX without its stop codon was used for Golden Gate assembly with pICH51266 [35S promoter+Ω promoter, Addgene no. 50267], pICSL50007 (3xFLAG, Addgene no. 50308) and pICH41432 (octopine synthase terminator, Addgene no. 50343) into binary vector pICH47732 (Addgene no. 48000).

To generate virus-induced gene silencing constructs, the silencing fragment was amplified from template cDNA of *NRCX* or *Prf* or TRV:NRC2/3, TRV:NRC4, TRV:NRC2/3/4 plasmid (Wu et al., 2016; Wu et al., 2017) by Phusion High-Fidelity DNA Polymerase (Thermo Fisher) using primers listed in *SI Appendix*, Table S3. The purified amplicons were directly used in Golden Gate assembly with pTRV-GG vector according to Duggan et al. (2021b).

To generate NRCX MADA motif chimera of NRC4 (MADA^NRCX^-NRC4^DV^), we followed a construction procedure of MADA^NRC2^-NRC4^DV^ and MADA^ZAR1^-NRC4^DV^ as described previously (Adachi et al., 2019b). The full-length sequence of *NRC4*^DV^ was amplified by Phusion High-Fidelity DNA Polymerase (Thermo Fisher) using primers listed in *SI Appendix*, Table S3. Purified amplicons were cloned into pCR8/GW/D-TOPO (Invitrogen) as a level 0 module. The level 0 plasmid was then used for Golden Gate assembly with pICH85281 [mannopine synthase promoter+Ω (MasΩpro), Addgene no. 50272], pICSL50009 (6xHA, Addgene no. 50309) and pICSL60008 [Arabidopsis heat shock protein terminator (HSPter), TSL SynBio] into the binary vector pICH47742 (Addgene no. 48001).

To generate MHD mutants of NRCX, the histidine (H) and aspartic acid (D) residues in the MHD motif were substituted by overlap extension PCR using Phusion High-Fidelity DNA Polymerase (Thermo Fisher). The pCR8::NRCX without its stop codon was used as a template. The mutagenesis primers are listed in *SI Appendix*, Table S3. The mutated NRCX was verified by DNA sequencing of the obtained plasmids.

To generate the RNAi construct, the silencing fragment was amplified from NRCX cDNA by Phusion High-Fidelity DNA Polymerase (Thermo Fisher Scientific) using primers listed in *SI Appendix*, Table S3. The purified amplicon was cloned into the pRNAi-GG vector (Yan et al., 2012).

Information of other constructs used for the cell death assays were described in *SI Appendix*, Table S4.

### Virus-induced gene silencing (VIGS)

VIGS was performed in *N. benthamiana* as previously described (Ratcliff et al., 2001). Suspensions of Agrobacterium Gv3101 strains carrying TRV RNA1 and TRV RNA2 were mixed in a 1:1 ratio in infiltration buffer (10 mM 2-[N-morpholine]-ethanesulfonic acid [MES]; 10 mM MgCl_2_; and 150 μM acetosyringone, pH 5.6) to a final OD_600_ of 0.25. Two-week-old *N. benthamiana* plants were infiltrated with the Agrobacterium suspensions for VIGS.

### Bioinformatic and phylogenetic analyses

Based on the NLR annotation and phylogenetic tree previously described in Harant et al. (2021), we extracted CC-NLR sequences of tomato, *N. benthamiana*, *A. thaliana* and sugar beet (*Beta vulgaris* ssp. *vulgaris* var. *altissima*). We then added NbS00004191g0008.1 and NbS00004611g0006.1, that lack the p-loop motif in the NB-ARC domain and are therefore missing in the previous CC-NLR list (Harant et al. 2021), and prepared a CC-NLR dataset (431 protein sequences, *Datasets*, File S1). Amino acid sequences of the CC-NLR dataset were aligned using MAFFT v.7 (Katoh and Standley, 2013). The gaps in the alignments were deleted manually and the NB-ARC domain sequences were used for generating phylogenetic tree (*Datasets*, File S2). The maximum likelihood tree based on the JTT model was made in RAxML version 8.2.12 (Stamatikis, 2014) and bootstrap values based on 100 iterations were shown in *Datasets*, File S3.

NRC-helper proteins were subjected to motif searches using MEME (Multiple EM for Motif Elicitation) v. 5.0.5 (Bailey and Elkan, 1994) with parameters ‘zero or one occurrence per sequence, top twenty motifs’, to detect consensus motifs conserved in > 80% of input sequences.

### Transient gene expression and cell death assays

Transient gene expression in *N. benthamiana* was performed by agroinfiltration according to methods described by Bos et al. (2006). Briefly, four-week-old *N. benthamiana* plants were infiltrated with Agrobacterium Gv3101 strains carrying the binary expression plasmids. The Agrobacterium suspensions were prepared in infiltration buffer (10 mM MES, 10 mM MgCl2, and 150 μM acetosyringone, pH5.6). To overexpress NRC MADA chimeras and MHD mutants, the concentration of each suspension was adjusted to OD_600_ = 0.5. To perform RNAi experiments in *N. benthamiana* leaves, we infiltrated Agrobacterium strains carrying RNAi constructs (OD_600_ = 0.5), together with different proteins described in *SI Appendix*, Table S4. Macroscopic cell death phenotypes were scored according to the scale of Segretin et al. (2014) modified to range from 0 (no visible necrosis) to 7 (fully confluent necrosis).

To stain dead cells by trypan blue, *N. benthamiana* leaves were transferred to a trypan blue solution (10 mL of lactic acid, 10 mL of glycerol, 10 g of phenol, 10 mL of water, and 10 mg of trypan blue) diluted in ethanol 1:1 and were incubated at 65 °C using a water bath for 1 hour. The leaves were then destained for 48 hours in a chloral hydrate solution (100 g of chloral hydrate, 5 mL of glycerol, and 30 mL of water).

### Protein immunoblotting

Protein samples were prepared from six discs (8 mm diameter) cut out of *N. benthamiana* leaves at 2 days after agroinfiltration and were homogenised in extraction buffer [10% (v/v) glycerol, 25 mM Tris-HCl, pH 7.5, 1 mM EDTA, 150 mM NaCl, 2% (w/v) PVPP, 10 mM DTT, 1x protease inhibitor cocktail (SIGMA), 0.5% (v/v) IGEPAL (SIGMA)]. The supernatant obtained after centrifugation at 12,000 x*g* for 10 min was used for SDS-PAGE. Immunoblotting was performed with HA-probe (F-7) HRP (Santa Cruz Biotech) in a 1:5,000 dilution. Equal loading was checked by taking images of the stained PVDF membranes with Pierce Reversible Protein Stain Kit (#24585, Thermo Fisher Scientific).

### RNA extraction and semi-quantitative RT-PCR

Total RNA was extracted using RNeasy Mini Kit (Qiagen). 500 ng RNA of each sample was subjected to reverse transcription using SuperScript® IV Reverse Transcriptase (Thermo Fisher Scientific). Semi-quantitative reverse transcription PCR (RT-PCR) was performed using DreamTaq (Thermo Fisher Scientific) with 25 to 30 amplification cycles followed by electrophoresis with 1.8% (w/v) agarose gel stained with Ethidium bromide. Primers used for RT-PCR are listed in *SI Appendix*, Table S3.

### RNA-seq analysis

Total RNAs of leaf, root and flower/bud tissue samples were extracted from six-week-old *N. benthamiana* plants using RNeasy Mini Kit (Qiagen). Three replicate each of the samples was sent for Illumina NovaSeq 6000 (40 M paired-end reads per sample, Novogene). Obtained RNA-seq reads were used to analyse expression profiles of NRC-helper genes in *N. benthamiana*. Reads were filtered and trimmed using FaQCs (Lo and Chain, 2014). The quality-trimmed reads were mapped to the reference *N. benthamiana* genome (Sol Genomics Network, v0.4.4) using HISAT2 (Kim et al., 2019). The number of read alignments in the gene regions were counted using featureCounts (Liao et al., 2014) and read counts were transformed into a Transcripts Per Million (TPM) value. Public RNA-seq reads from six-week-old *N. bethamiana* leaves with or without *Pseudomonas fluorescens* inoculation (Pombo et al., 2019), were also analysed as described above (Accession Numbers: SRP118889).

### Accession numbers

NRC sequences used in this study can be found in the GenBank/EMBL and Solanaceae Genomics Network (SGN) databases with the following accession numbers: NbNRC2 (NbS00018282g0019.1 and NbS00026706g0016.1), NbNRC3 (MK692736.1), NbNRC4 (MK692737) and NbNRCX (NbS00030243g0001.1).

## ACKNOWLEDGMENTS

We are thankful to Jiorgos Kourelis, Lida Derevnina and colleagues for valuable discussions and ideas. We thank Mark Youles of TSL SynBio for invaluable technical support. This work was funded by the Gatsby Charitable Foundation and Biotechnology and Biological Sciences Research Council (BBSRC, UK). S.K. also receives funding from the European Research Council (ERC BLASTOFF projects). H.A. was funded by the Japan Society for the Promotion of Science (JSPS).

## AUTHOR CONTRIBUTIONS

Conceptualization: H.A., C.H.W., S.K.; Data curation: H.A., T.S., C.H.W.; Formal analysis: H.A., T.S.; Investigation: H.A., T.S., C.H.W.; Methodology: H.A., T.S., C.H.W.; Resources: H.A., T.S., A.H., C.D., T.O.B., C.H.W.; Supervision: H.A., C.H.W, S.K.; Funding acquisition: H.A., S.K.; Project administration: S.K.; Writing initial draft: H.A., S.K.; Editing: H.A., T.S., C.D., T.O.B., C.H.W., S.K.

---

**Fig. S1.**
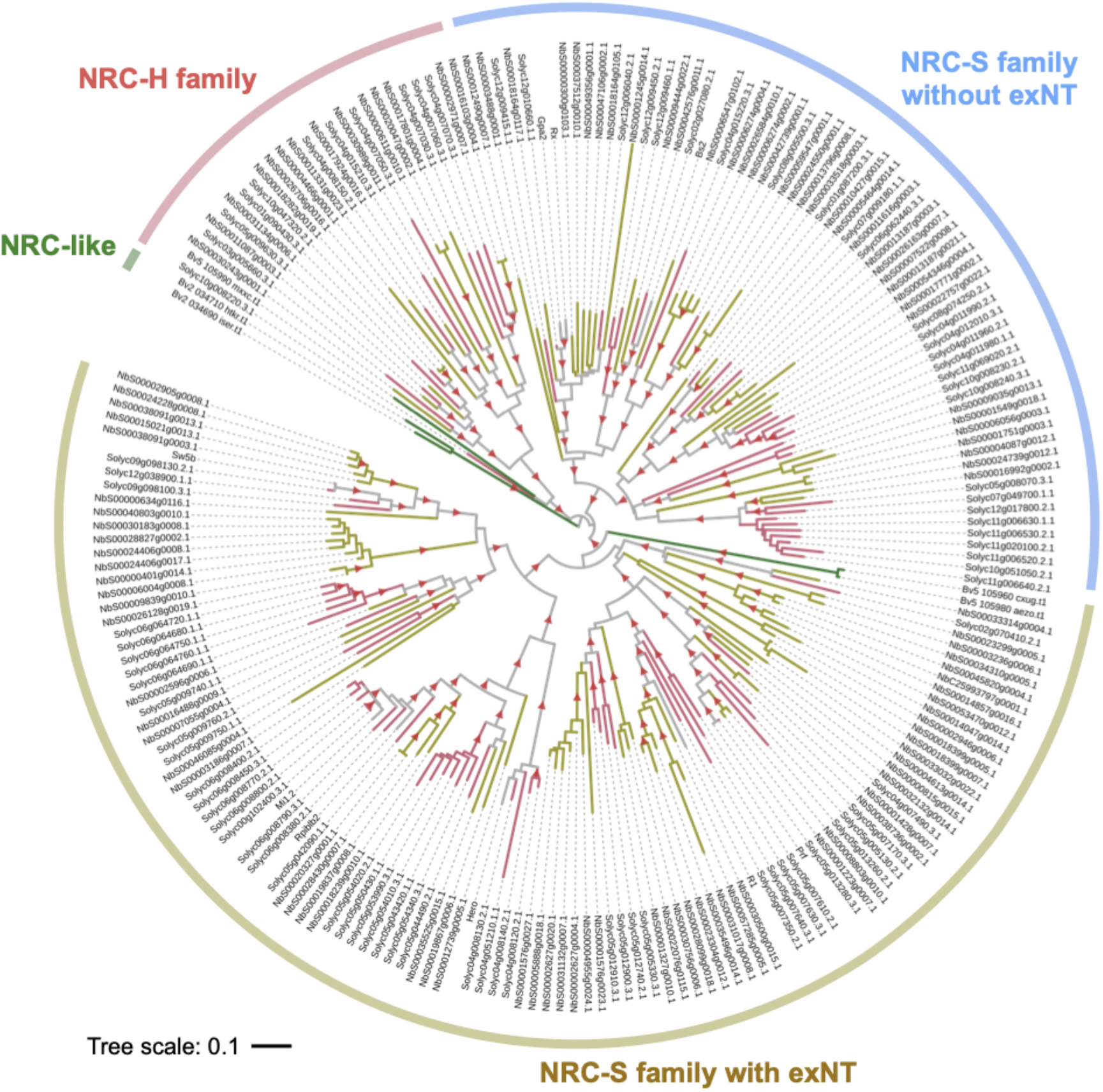
Phylogenetic tree of NRC superclade. NRC-sensor (NRC-S) and NRC-helper (NRC-H) proteins identified in Adachi et al. (2019b) were used for the MAFFT multiple alignment and phylogenetic analyses. The phylogenetic tree was constructed with the NB-ARC domain sequences in MEGA7 by the neighbour-joining method. Each leaf is labelled with different colour ranges indicating plant species, *N. benthamiana* (NbS-), tomato (Solyc-) and sugar beet (Bv-). The NRC-S clade is divided into NLRs that lack an extended N-terminal domain (exNT) prior to their CC domain and those that carry an exNT. Red arrow heads indicate bootstrap support > 0.7. The scale bars indicate the evolutionary distance in amino acid substitution per site.

**Fig. S2.**
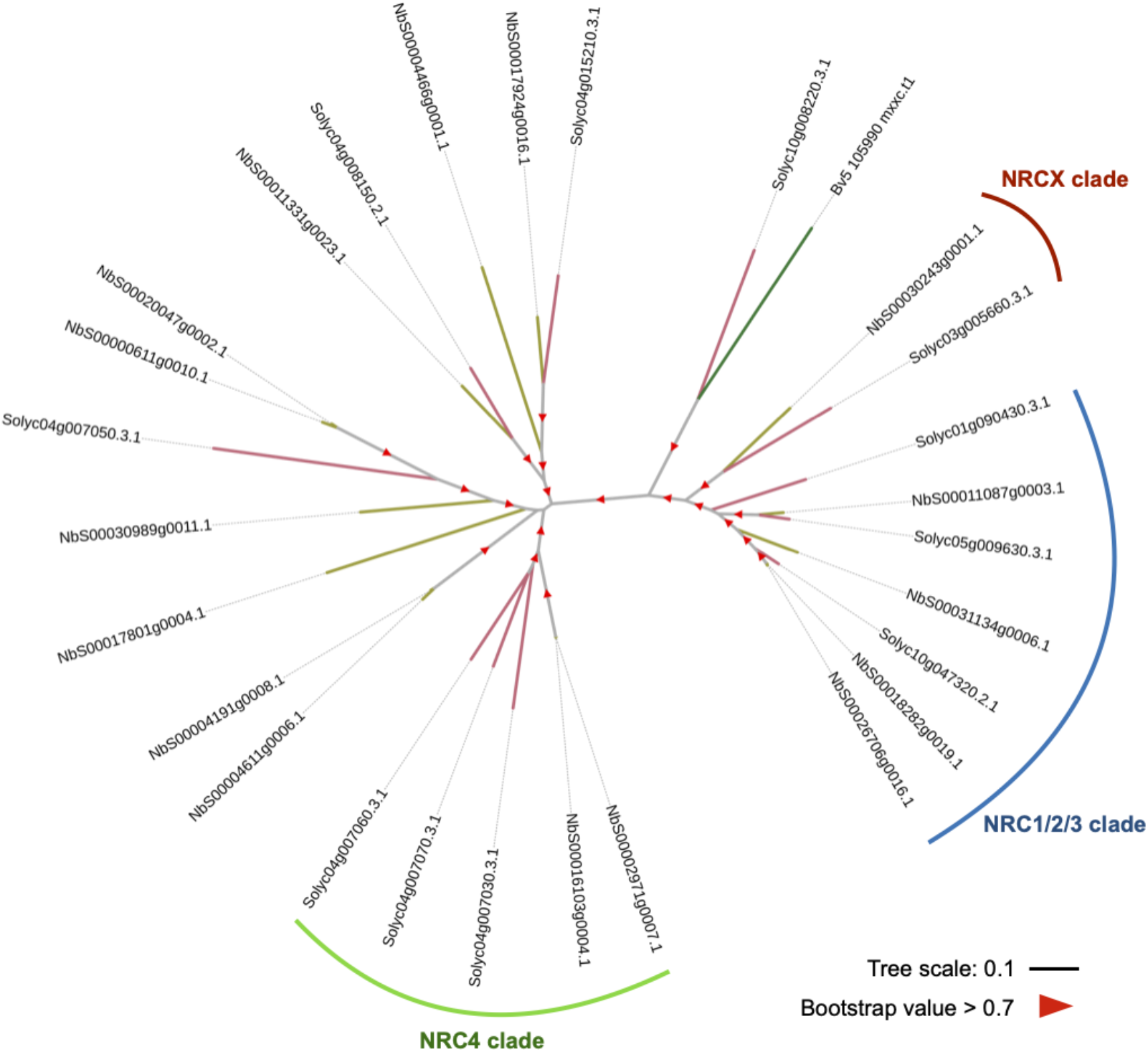
NRC-H subclade shown in Figure 2B.

**Fig. S3.**
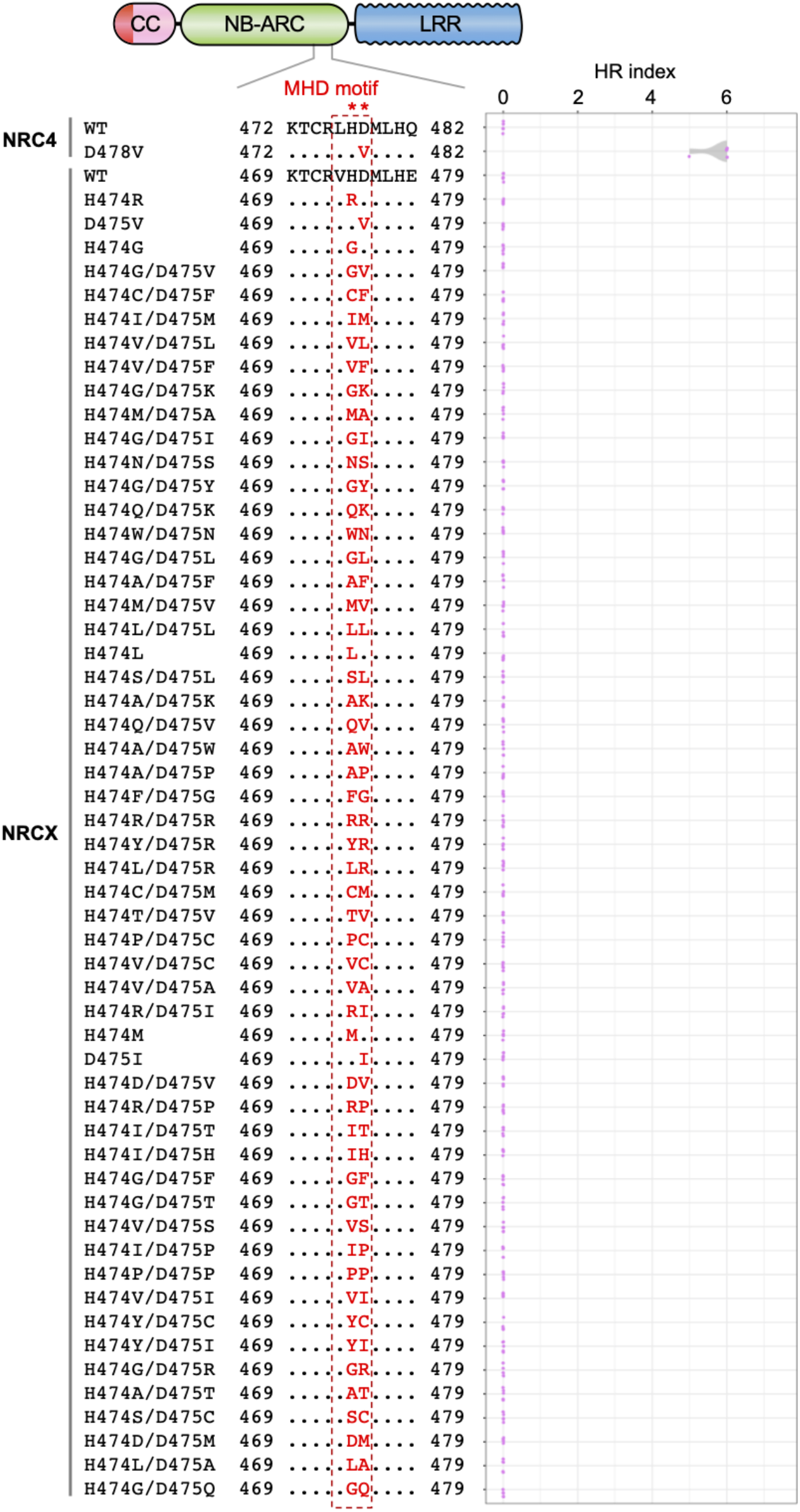
55 independent NRCX MHD mutants do not cause autoactive cell death in *N. benthamiana*. Cell death phenotypes were scored at an HR index at 5 days after agroinfiltration to express NRC4^WT^, NRCX^WT^ and the MHD mutants in *N. benthamiana* leaves. Quantification data are from 5 independent biological replicates.

**Fig. S4.**
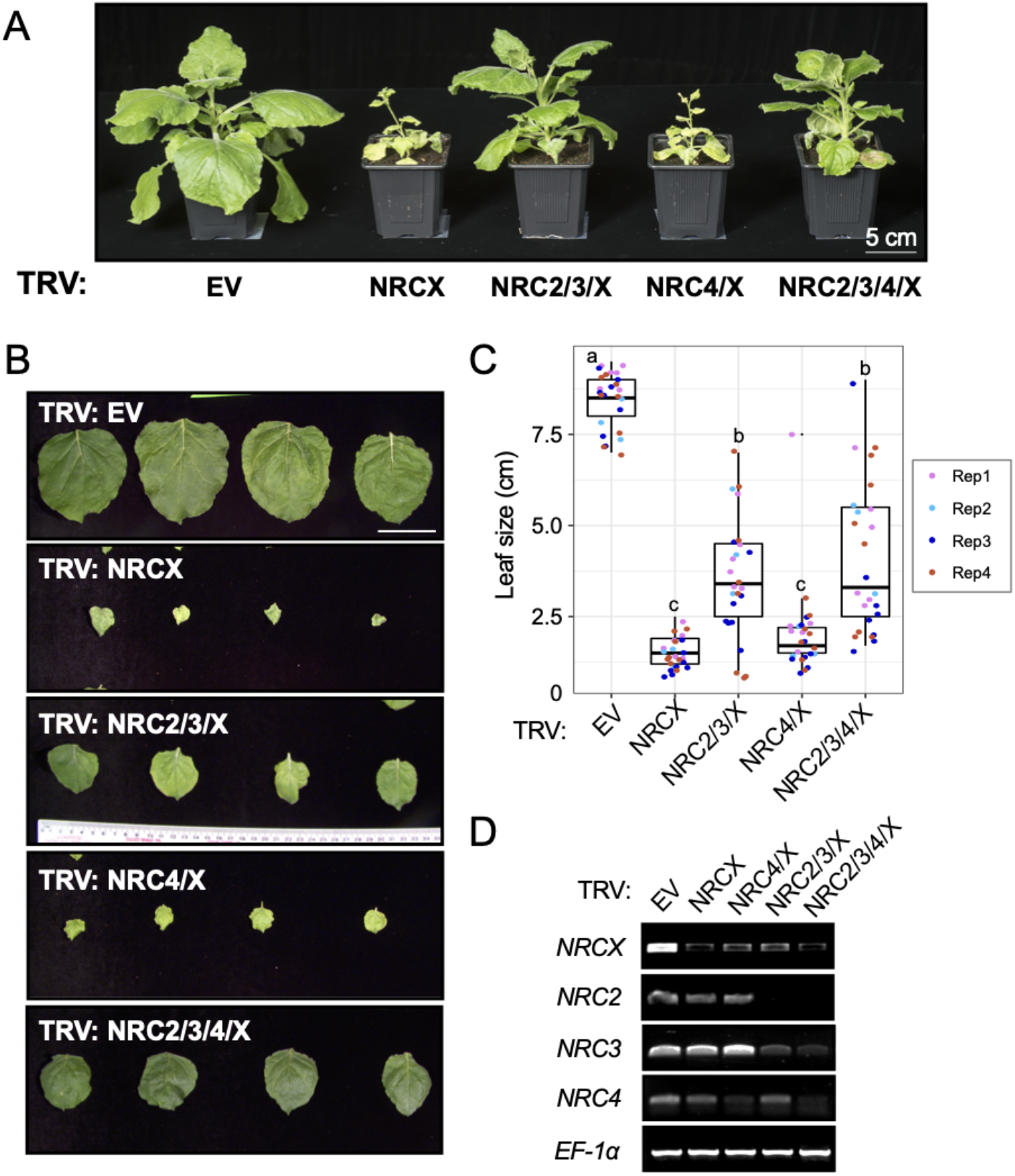
Co-silencing of *NRC2* and *NRC3* partially suppresses TRV:NRCX dwarf phenotype in *N. benthamiana*. **(A)** The morphology of 6-week-old *NRCX-*, *NRC2/3/X-*, *NRC4/X-* and *NRC2/3/4/X-*silenced *N. benthamiana* plants. 2-week-old *N. benthamiana* plants were infiltrated with *Agrobacterium* strains carrying VIGS constructs, and photographs were taken 4 weeks after the agroinfiltration. TRV empty vector (TRV:EV) was used as a negative control. **(B, C)** Quantification of the leaf size. One leaf per each plant was harvested from the same position (the 5th leaf from cotyledons) and was used for measuring the leaf diameter. Statistical differences among the samples were analyzed with Tukey’s HSD test (p<0.01). Scale bars = 5 cm. (D) Specific gene silencing of *NRCX* or multiple *NRC* genes in TRV:NRC-infected plants. Leaf samples were collected for RNA extraction at 3 weeks after agroinfiltration expressing VIGS constructs. The expression of *NRCX* and other *NRC* genes were analyzed in semi-quantitative RT-PCR using specific primer sets. Elongation factor 1α (EF-1α) was used as an internal control. Scale bars = 5 cm.

**Fig. S5.**
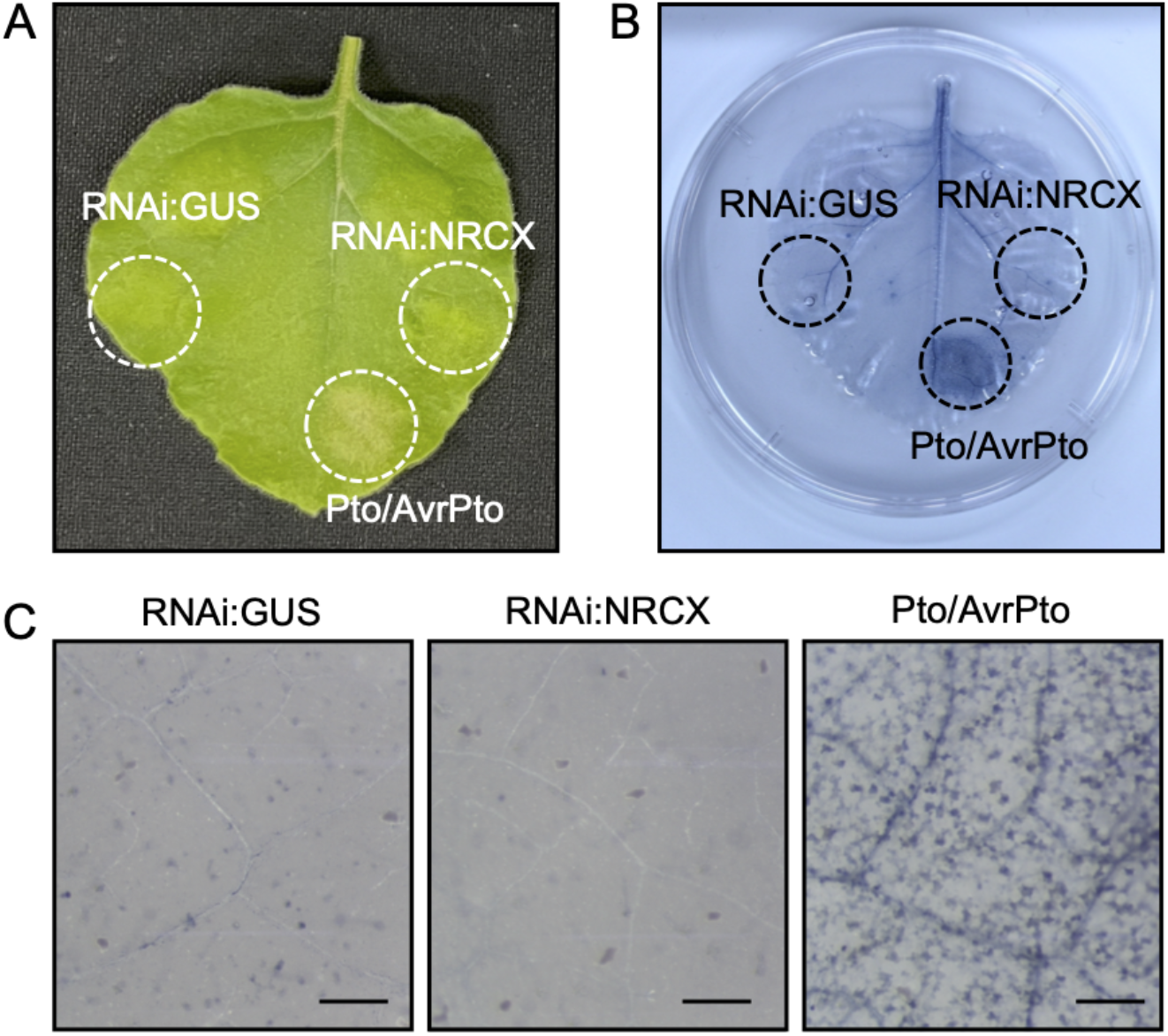
RNA interference of *NRCX* does not cause cell death in *N. benthamiana* leaves. **(A)** Macroscopic cell death phenotype after expressing RNAi:GUS, RNAi:NRCX or Pto/AvrPto by agroinfiltration. Photograph was taken at 5 days after the agroinfiltration. **(B)** Cell death was detected by trypan blue staining at 5 days after the agroinfiltration. **(C)** Microscopic cell death phenotype. Dead cells were stained by trypan blue. Images describe representative data of 8 replicates from 2 independent experiments. Scale bars are 300 μm.

**Fig. S6.**
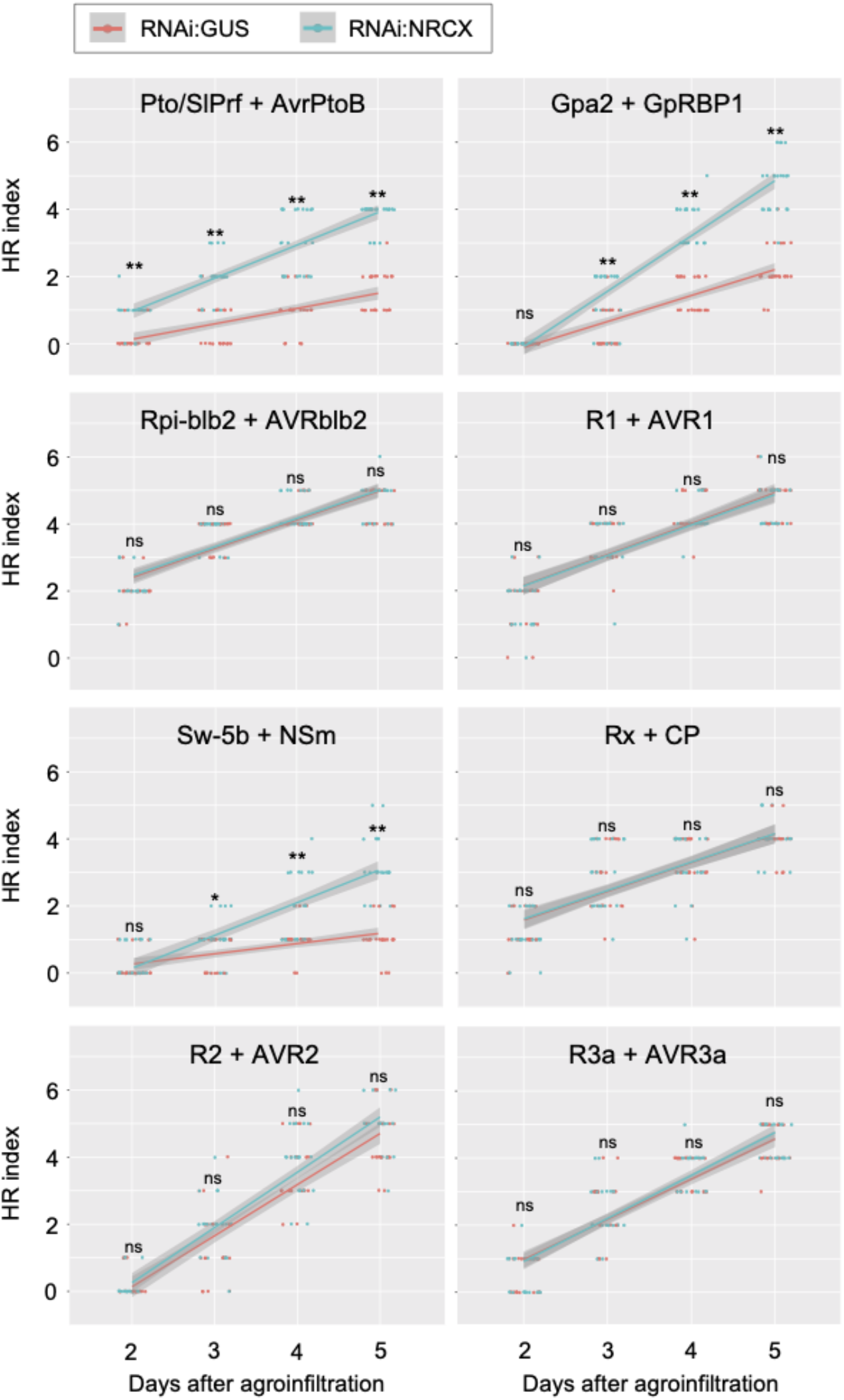
Time-lapse quantification of NRC-S/AVR-triggered hypersensitive cell death in *NRCX* silenced leaves. Cell death intensity was scored at 2-5 days after the agroinfiltration as described in Fig 6. The HR index plots are based on three independent experiments. Asterisks indicate statistically significant differences with *t* test (*p<0.05 and **p<0.01). Pink and blue line plots indicate mean values of RNAi:GUS and RNAi:NRCX samples at each time point.

**Fig. S7.**
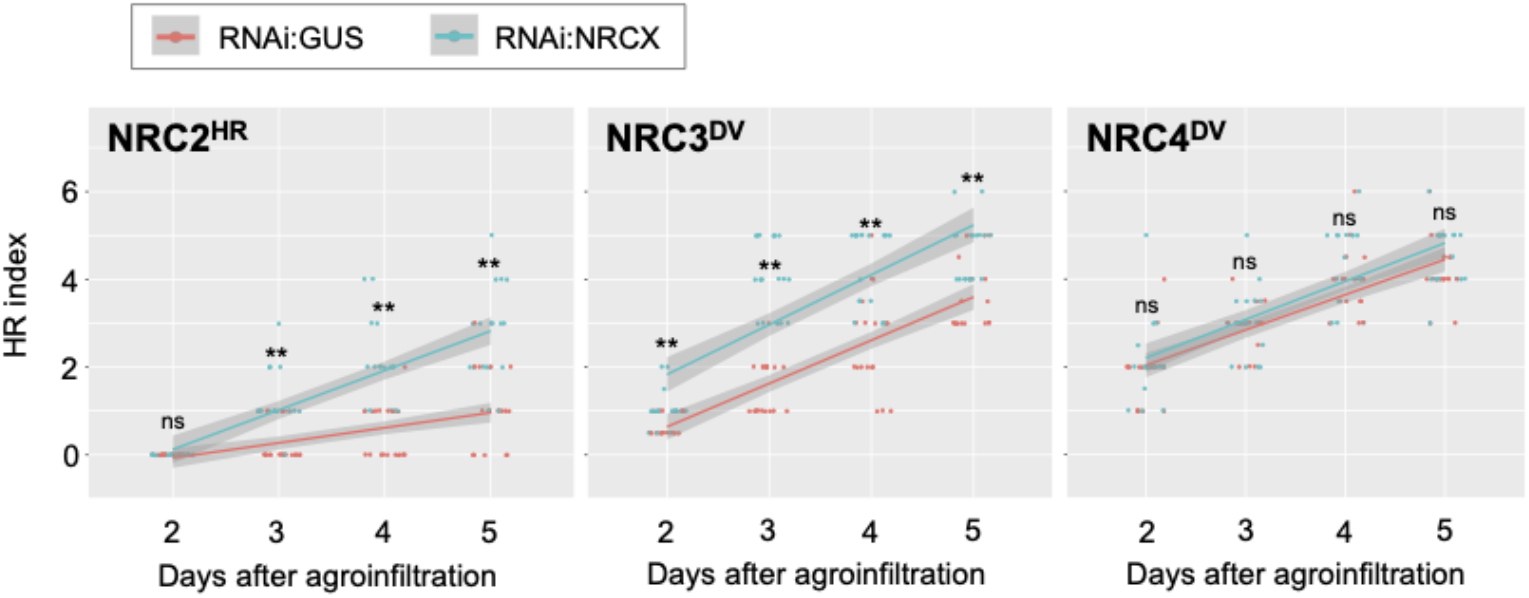
Time-lapse quantification of NRC-H autoactive cell death in *NRCX* silenced leaves. Cell death intensity was scored at 2-5 days after the agroinfiltration as described in Fig 6. The HR index plots are based on three independent experiments. Asterisks indicate statistically significant differences with *t* test (**p<0.01). Pink and blue line plots indicate mean values of RNAi:GUS and RNAi:NRCX samples at each time point.

**Fig. S8.**
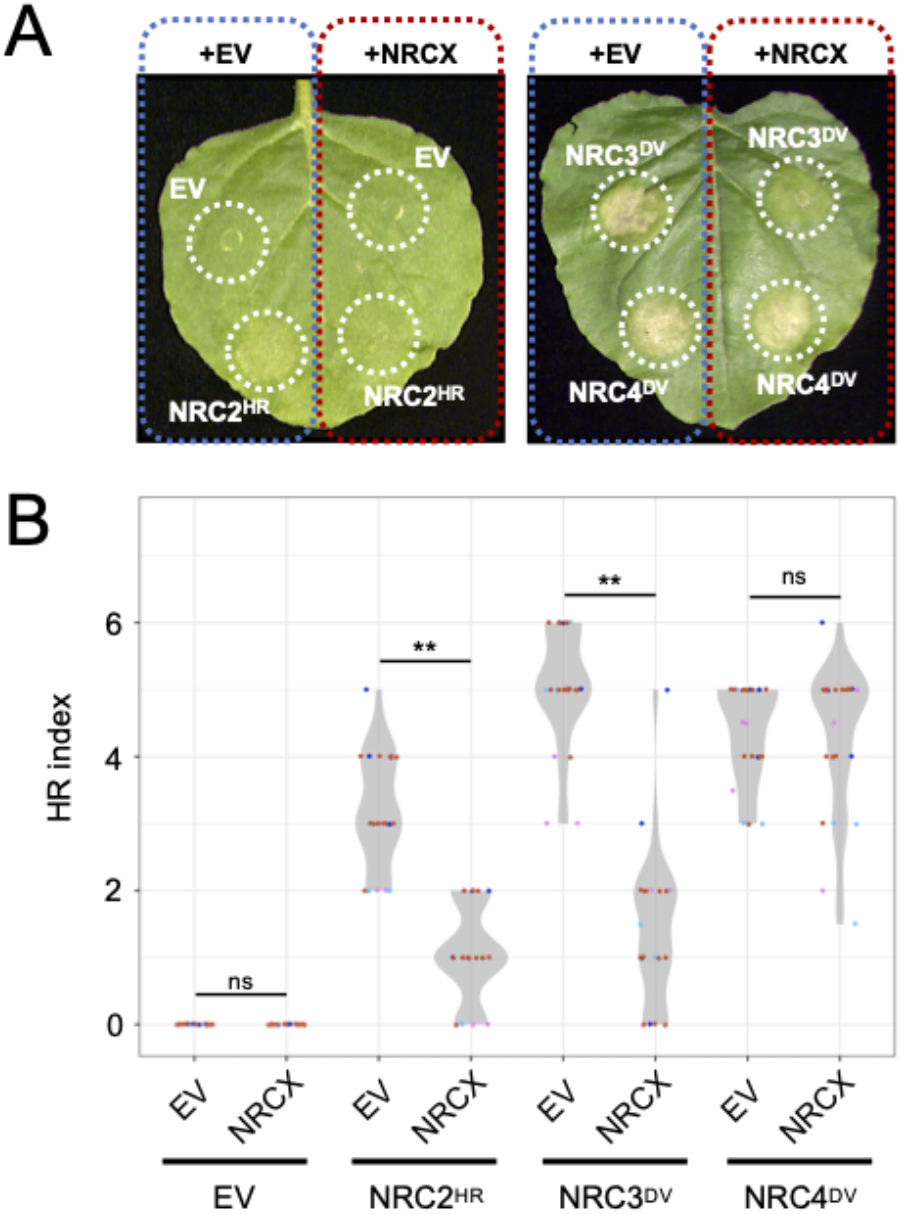
Overexpression of wild-type NRCX compromises autoactive cell death of NRC2 and NRC3, but not NRC4. **(A)** Photo of representative *N. benthamiana* leaves showing autoactive cell death after co-expression of empty vector (EV; control) and wild-type NRCX with NRC2^HR^, NRC3^DV^ and NRC4^DV^. Photographs were taken at 5 days after agroinfiltration. **(B)** Violin plots showing cell death intensity scored as an HR index at 5 days after the agroinfiltration. The HR index plots are based on four independent experiments. Asterisks indicate statistically significant differences with *t* test (**p<0.01).

## Supplemental data

**Table S1. Transcriptome profiles of *Nicotiana benthamiana* NLR genes.**

**Table S2. Expression ratios of *NRC2*, *NRC3* and *NRC4* compared to *NRCX*.**

**Table S3. Primers used in this study.**

**Table S4. List of NLR and corresponding AVR effector used in cell death assays.**

**File S1. Amino acid sequences of full-length CC-NLRs used for phylogenetic analysis in Figure 2A.**

**File S2. Amino acid sequences for CC-NLR phylogenetic tree in Figure 2A.**

**File S3. CC-NLR phylogenetic tree file in Figure 2A.**

