## Supplementary material for "An atypical NLR protein modulates the NRC immune receptor network": Table S2_expression ratio.pptx

### Slide 1
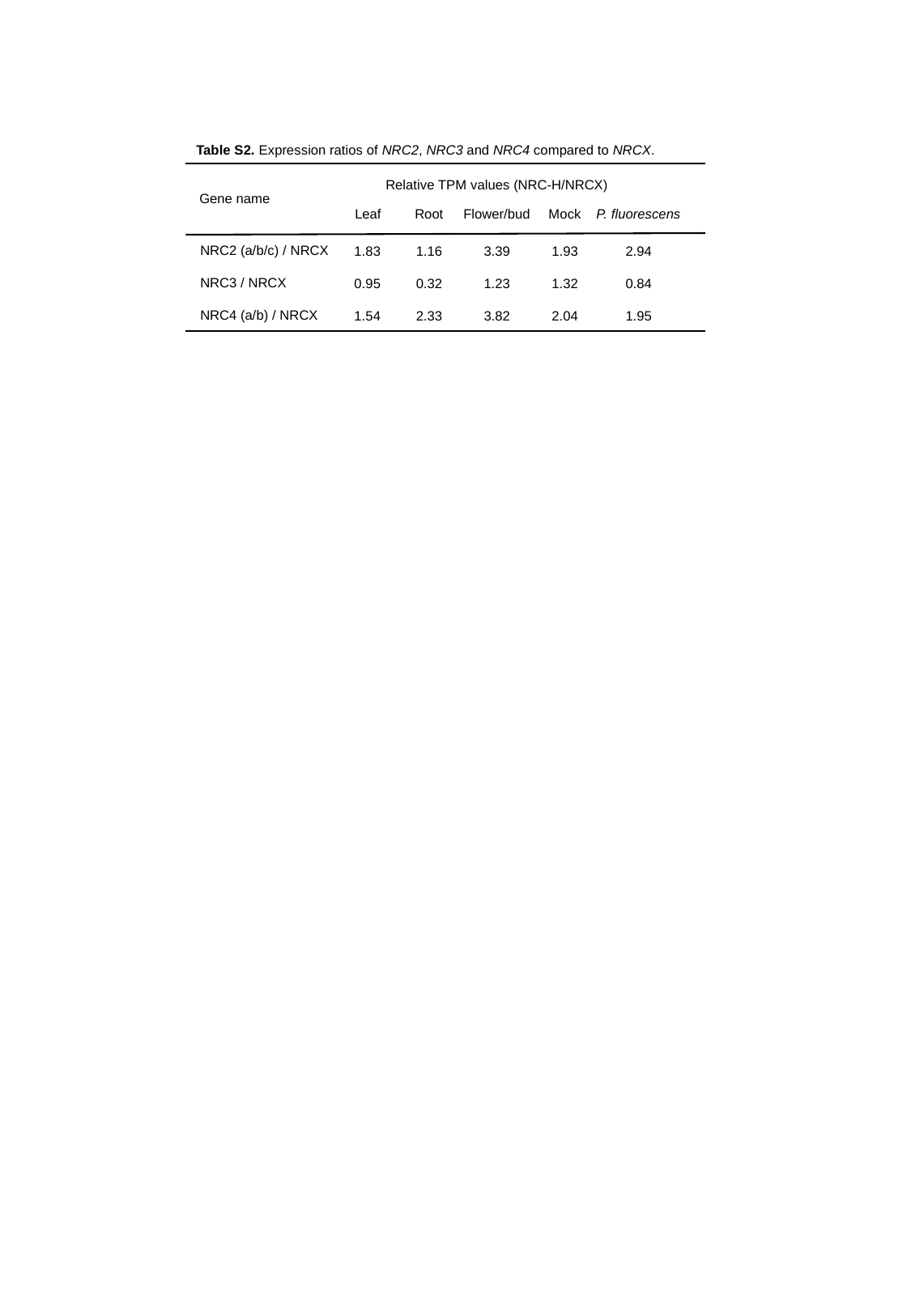

Table S2. Expression ratios of NRC2, NRC3 and NRC4 compared to NRCX.
Relative TPM values (NRC-H/NRCX)
Gene name
Leaf
Root
Flower/bud
Mock
P. fluorescens
NRC2 (a/b/c) / NRCX
NRC3 / NRCX
NRC4 (a/b) / NRCX
1.83
0.95
1.54
1.16
0.32
2.33
3.39
1.23
3.82
1.93
1.32
2.04
2.94
0.84
1.95
