## Supplementary material for "An atypical NLR protein modulates the NRC immune receptor network": Table S3_primers.pptx

### Slide 1
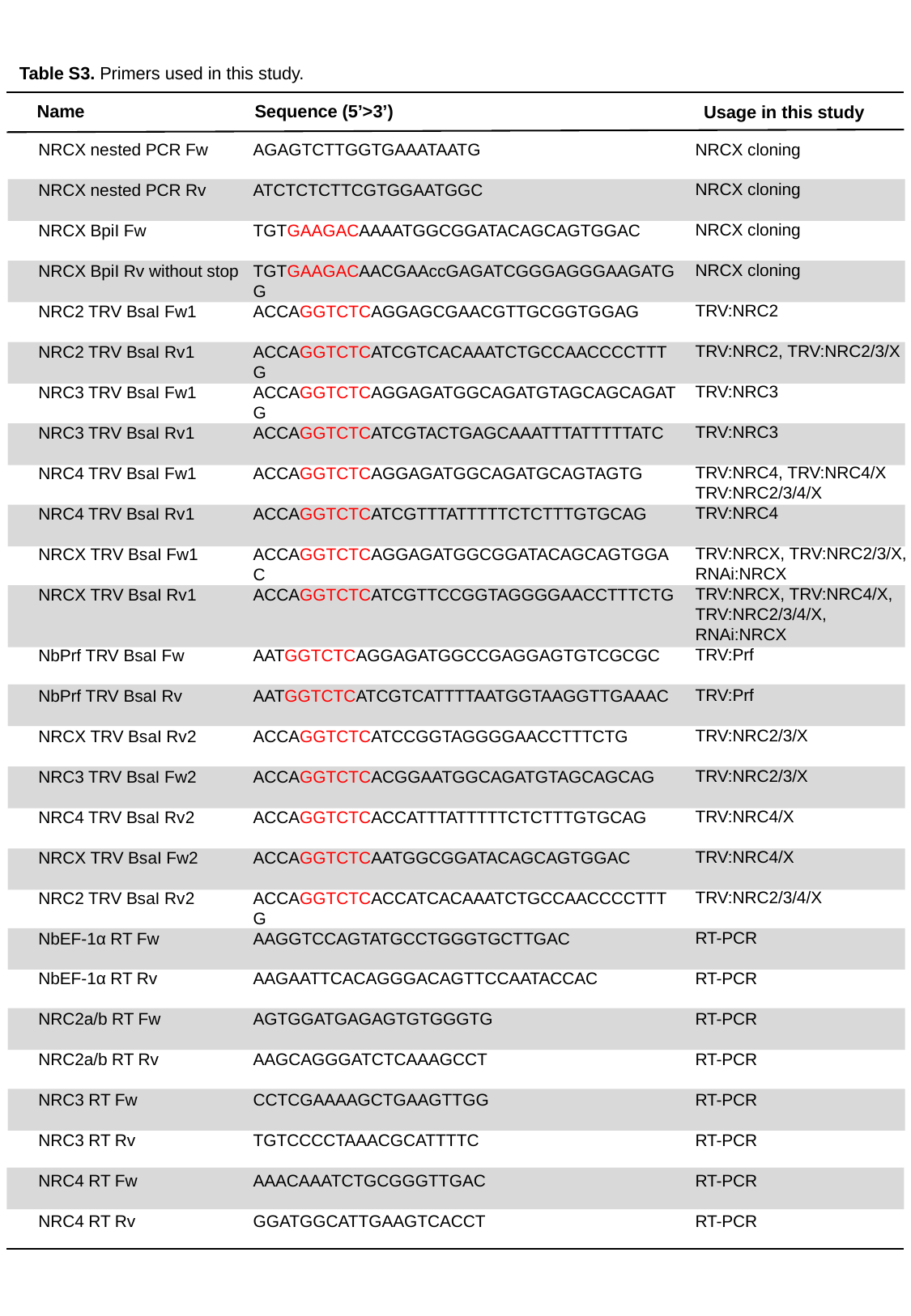

Table S3. Primers used in this study.
Name
Sequence (5’>3’)
Usage in this study
NRCX cloning
NRCX cloning
NRCX cloning
NRCX cloning
TRV:NRC2
TRV:NRC2, TRV:NRC2/3/X
TRV:NRC3
TRV:NRC3
TRV:NRC4, TRV:NRC4/X
TRV:NRC2/3/4/X
TRV:NRC4
TRV:NRCX, TRV:NRC2/3/X, RNAi:NRCX
TRV:NRCX, TRV:NRC4/X, TRV:NRC2/3/4/X, RNAi:NRCX
TRV:Prf
TRV:Prf
TRV:NRC2/3/X
TRV:NRC2/3/X
TRV:NRC4/X
TRV:NRC4/X
TRV:NRC2/3/4/X
RT-PCR
RT-PCR
RT-PCR
RT-PCR
RT-PCR
RT-PCR
RT-PCR
RT-PCR
NRCX nested PCR Fw
NRCX nested PCR Rv
NRCX BpiI Fw
NRCX BpiI Rv without stop
NRC2 TRV BsaI Fw1
NRC2 TRV BsaI Rv1
NRC3 TRV BsaI Fw1
NRC3 TRV BsaI Rv1
NRC4 TRV BsaI Fw1
NRC4 TRV BsaI Rv1
NRCX TRV BsaI Fw1
NRCX TRV BsaI Rv1
NbPrf TRV BsaI Fw
NbPrf TRV BsaI Rv
NRCX TRV BsaI Rv2
NRC3 TRV BsaI Fw2
NRC4 TRV BsaI Rv2
NRCX TRV BsaI Fw2
NRC2 TRV BsaI Rv2
NbEF-1α RT Fw
NbEF-1α RT Rv
NRC2a/b RT Fw
NRC2a/b RT Rv
NRC3 RT Fw
NRC3 RT Rv
NRC4 RT Fw
NRC4 RT Rv
AGAGTCTTGGTGAAATAATG
ATCTCTCTTCGTGGAATGGC
TGTGAAGACAAAATGGCGGATACAGCAGTGGAC
TGTGAAGACAACGAAccGAGATCGGGAGGGAAGATGG
ACCAGGTCTCAGGAGCGAACGTTGCGGTGGAG
ACCAGGTCTCATCGTCACAAATCTGCCAACCCCTTTG
ACCAGGTCTCAGGAGATGGCAGATGTAGCAGCAGATG
ACCAGGTCTCATCGTACTGAGCAAATTTATTTTTATC
ACCAGGTCTCAGGAGATGGCAGATGCAGTAGTG
ACCAGGTCTCATCGTTTATTTTTCTCTTTGTGCAG
ACCAGGTCTCAGGAGATGGCGGATACAGCAGTGGAC
ACCAGGTCTCATCGTTCCGGTAGGGGAACCTTTCTG
AATGGTCTCAGGAGATGGCCGAGGAGTGTCGCGC
AATGGTCTCATCGTCATTTTAATGGTAAGGTTGAAAC
ACCAGGTCTCATCCGGTAGGGGAACCTTTCTG
ACCAGGTCTCACGGAATGGCAGATGTAGCAGCAG
ACCAGGTCTCACCATTTATTTTTCTCTTTGTGCAG
ACCAGGTCTCAATGGCGGATACAGCAGTGGAC
ACCAGGTCTCACCATCACAAATCTGCCAACCCCTTTG
AAGGTCCAGTATGCCTGGGTGCTTGAC
AAGAATTCACAGGGACAGTTCCAATACCAC
AGTGGATGAGAGTGTGGGTG
AAGCAGGGATCTCAAAGCCT
CCTCGAAAAGCTGAAGTTGG
TGTCCCCTAAACGCATTTTC
AAACAAATCTGCGGGTTGAC
GGATGGCATTGAAGTCACCT

### Slide 2
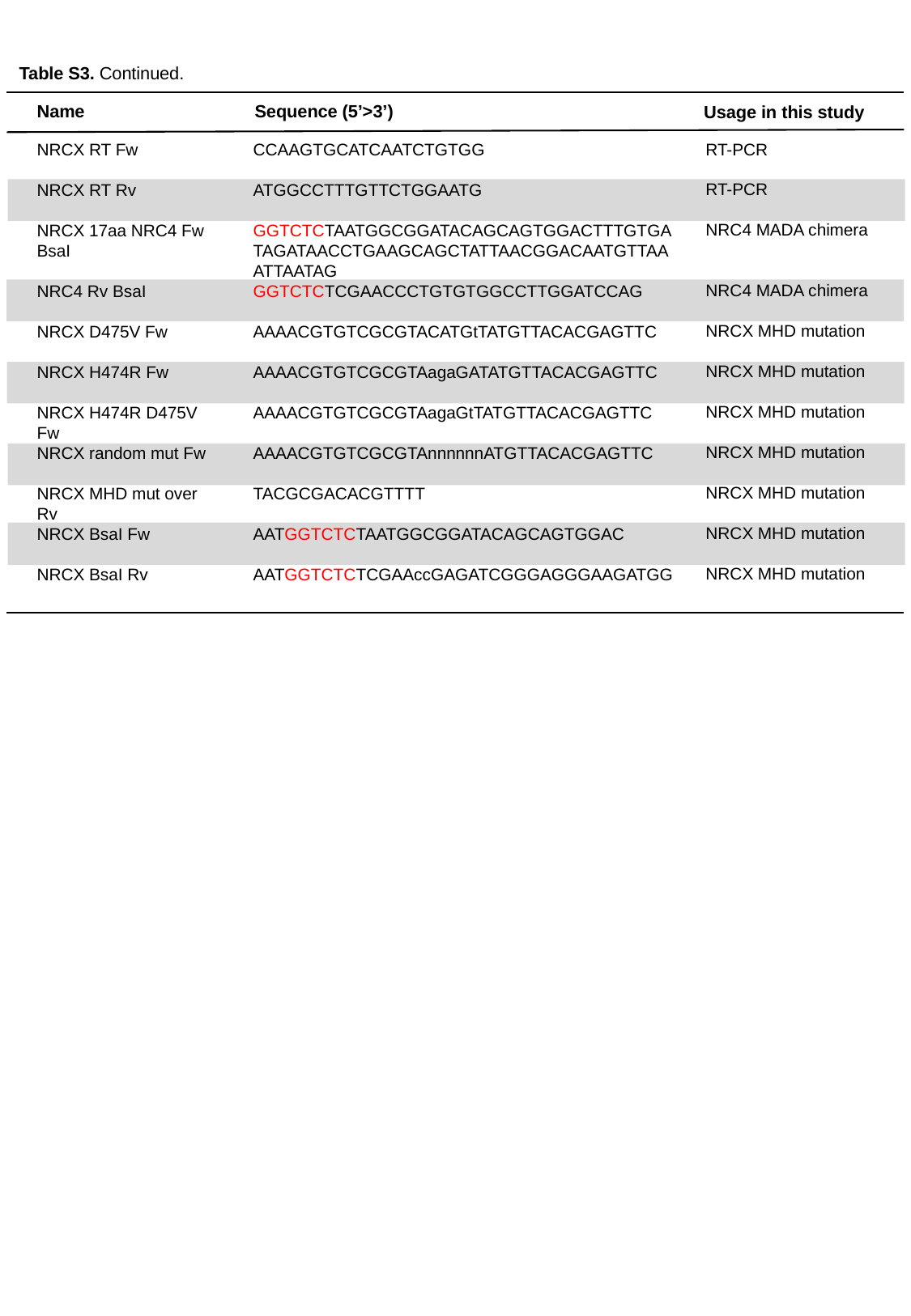

Table S3. Continued.
Name
Sequence (5’>3’)
Usage in this study
RT-PCR
RT-PCR
NRC4 MADA chimera
NRC4 MADA chimera
NRCX MHD mutation
NRCX MHD mutation
NRCX MHD mutation
NRCX MHD mutation
NRCX MHD mutation
NRCX MHD mutation
NRCX MHD mutation
NRCX RT Fw
NRCX RT Rv
NRCX 17aa NRC4 Fw BsaI
NRC4 Rv BsaI
NRCX D475V Fw
NRCX H474R Fw
NRCX H474R D475V Fw
NRCX random mut Fw
NRCX MHD mut over Rv
NRCX BsaI Fw
NRCX BsaI Rv
CCAAGTGCATCAATCTGTGG
ATGGCCTTTGTTCTGGAATG
GGTCTCTAATGGCGGATACAGCAGTGGACTTTGTGATAGATAACCTGAAGCAGCTATTAACGGACAATGTTAAATTAATAG
GGTCTCTCGAACCCTGTGTGGCCTTGGATCCAG
AAAACGTGTCGCGTACATGtTATGTTACACGAGTTC
AAAACGTGTCGCGTAagaGATATGTTACACGAGTTC
AAAACGTGTCGCGTAagaGtTATGTTACACGAGTTC
AAAACGTGTCGCGTAnnnnnnATGTTACACGAGTTC
TACGCGACACGTTTT
AATGGTCTCTAATGGCGGATACAGCAGTGGAC
AATGGTCTCTCGAAccGAGATCGGGAGGGAAGATGG
