## Supplementary material for "An atypical NLR protein modulates the NRC immune receptor network": Table S4_NLR_AVR.pptx

### Slide 1
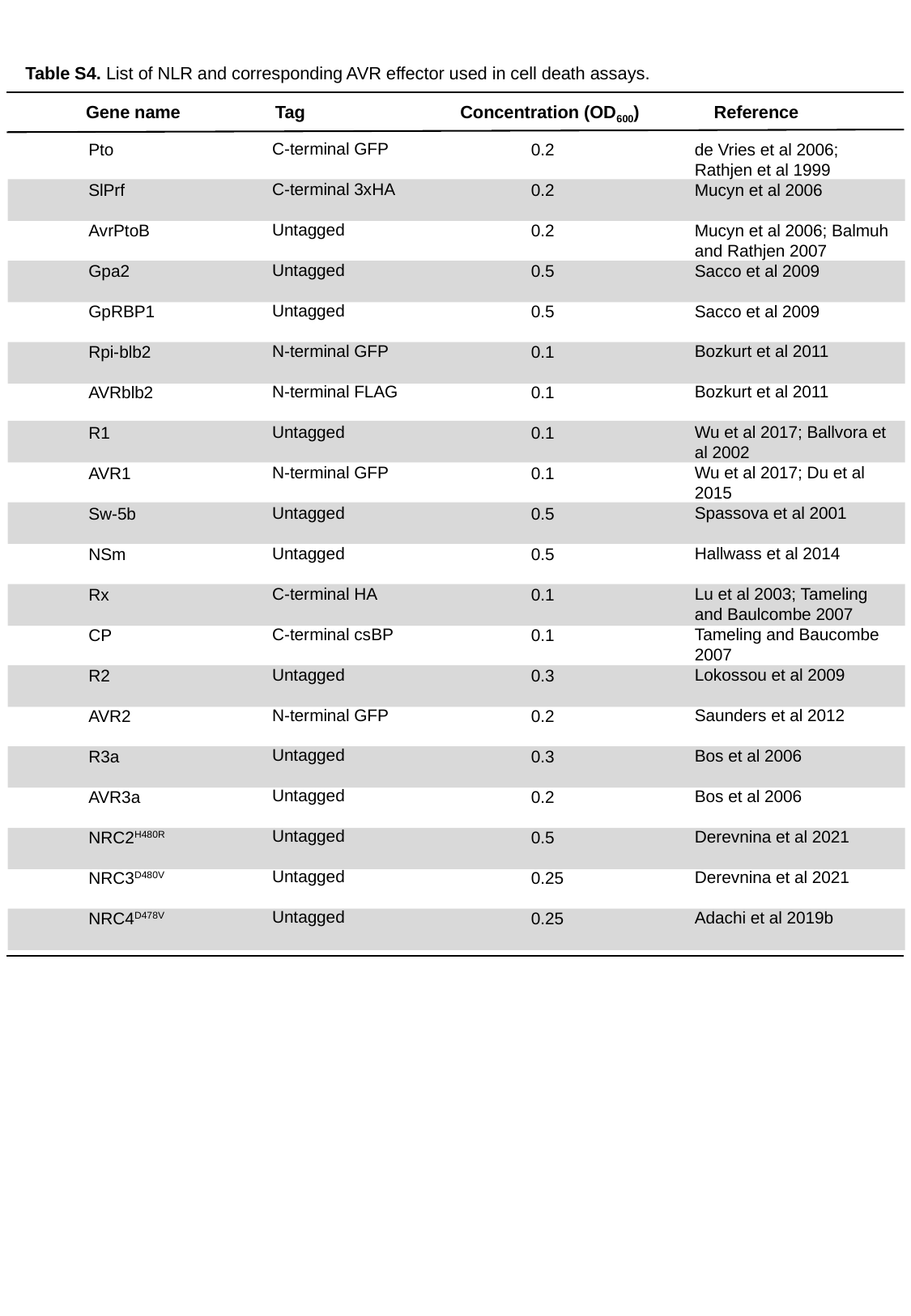

Table S4. List of NLR and corresponding AVR effector used in cell death assays.
Gene name
Tag
Concentration (OD_600_)
Reference
C-terminal GFP
C-terminal 3xHA
Untagged
Untagged
Untagged
N-terminal GFP
N-terminal FLAG
Untagged
N-terminal GFP
Untagged
Untagged
C-terminal HA
C-terminal csBP
Untagged
N-terminal GFP
Untagged
Untagged
Untagged
Untagged
Untagged
de Vries et al 2006; Rathjen et al 1999
Mucyn et al 2006
Mucyn et al 2006; Balmuh and Rathjen 2007
Sacco et al 2009
Sacco et al 2009
Bozkurt et al 2011
Bozkurt et al 2011
Wu et al 2017; Ballvora et al 2002
Wu et al 2017; Du et al 2015
Spassova et al 2001
Hallwass et al 2014
Lu et al 2003; Tameling and Baulcombe 2007
Tameling and Baucombe 2007
Lokossou et al 2009
Saunders et al 2012
Bos et al 2006
Bos et al 2006
Derevnina et al 2021
Derevnina et al 2021
Adachi et al 2019b
Pto
SlPrf
AvrPtoB
Gpa2
GpRBP1
Rpi-blb2
AVRblb2
R1
AVR1
Sw-5b
NSm
Rx
CP
R2
AVR2
R3a
AVR3a
NRC2^H480R^
NRC3^D480V^
NRC4^D478V^
0.2
0.2
0.2
0.5
0.5
0.1
0.1
0.1
0.1
0.5
0.5
0.1
0.1
0.3
0.2
0.3
0.2
0.5
0.25
0.25
